# Upregulated NF-κB pathway proteins may underlie *APOE44* associated astrocyte phenotypes in sporadic Alzheimer’s disease

**DOI:** 10.1101/2023.04.19.537428

**Authors:** Adele Pryce Roberts, Karolina Dec, Branduff McAllister, Victoria Tyrrell, Valerie B O’Donnell, Adrian Harwood, Julie Williams

**Affiliations:** UK Dementia Research Institute at Cardiff, Cardiff University, Hadyn Ellis Building, Maindy Road, CF24 4HQ, Cardiff, UK; Cardiff School of Biosciences, Cardiff University, Hadyn Ellis Building, Maindy Road, CF24 4HQ, Cardiff, UK; Division of Psychological Medicine and Clinical Neurosciences, Cardiff University, Cardiff, UK; Systems Immunity Research Institute and Division of Infection and Immunity, School of Medicine, Cardiff University, CF14 4XN, Cardiff, UK

**Keywords:** Alzheimer’s, *APOE*, Apolipoprotein E, CRISPR, astrocytes, NF-κB

## Abstract

The Apolipoprotein-E4 allele (APOE) is the strongest genetic risk factor for sporadic Alzheimer’s disease but its role in disease pathogenesis is incompletely understood. The *APOE* gene encodes Apolipoprotein E (ApoE). Astrocytes are the main source of ApoE in the central nervous system (CNS) and are essential for homeostasis in health and disease. In response to CNS insult, a coordinated multicellular inflammatory response is triggered causing reactive astrogliosis with changes in astrocytic gene expression, cellular structure and function.

Human embryonic stem-cells with the ‘neutral’ *APOE33* genotype were edited using CRISPR Cas-9 gene-editing to create isogenic *APOE* lines with an APOE44 genotype. Quiescent astrocytes were differentiated then stimulated with TNF-α, IL1α and C1q inducing an astrogliotic A1 phenotype. Several potentially pathological *APOE44*-related phenotypes were identified in both quiescent cells and reactive A1 astrocytes including significantly decreased phagocytosis, impaired glutamate and a defective immunomodulatory response.

In quiescent *APOE44* astrocytes there was significantly decreased secretion of IL6, IL8 and several oxylipins. In A1 astrocytes there was a pro-inflammatory phenotype in APOE44 astrocytes with increases in GRO, ENA78, IL6 and IL8, a decrease in IL10 as well as significant differences in oxylipin expression. As TNF-α induced signaling in astrocytes is driven by Nuclear factor kappa B (NF-κB) proteins of this pathway were measured. Significantly higher levels of the p50, p65 and IκBα sub-units were found in both quiescent and A1 *APOE44* astrocytes. This suggests that perturbation of NF-κB signaling may contribute to the damaging *APOE44* cell phenotypes observed providing a new direction for targeted disease therapeutics.

## 1. Introduction

The Apolipoprotein-E4 allele (*APOE*) is the strongest genetic risk factor for sporadic Alzheimer’s disease (sAD). Compared to non-carriers of *APOE4*, the increased risk of sAD is 9–15 fold in APOE4 homozygotes (1) equating to a lifetime risk of developing sAD of over 50% (2). Despite its importance, the role of *APOE* in disease pathogenesis is incompletely understood. *APOE* encodes apolipoprotein E (ApoE), a protein with a large number of functions both within and outside the central nervous system (CNS). In the brain, ApoE protein is mainly produced by astrocytes (3). Having once been considered neuronal scaffolding, it is now clear that astrocytes are essential to central nervous system (CNS) homeostasis (4).

In addition to their many physiological functions in the healthy CNS, astrocytes are also essential when the brain is compromised. In conjunction with microglia, they are the foundation of the CNS’s innate immune system (5) which is consistently implicated in sAD by pathway analyses (6). In response to damage and disease a coordinated multicellular, inflammatory response is triggered (7) causing reactive astrogliosis (8) with changes in astrocytic gene expression, cellular structure and function (9). Reactive astrocytes were first described by Alois Alzheimer as one of the pathological hallmarks of sAD but are relatively under studied in comparison to amyloid plaques and neurofibrillary tangles (NFTs) (9). Astrogliosis has essential beneficial functions but may also lead to harmful effects (8).

A1 astrocytes are a subtype of reactive astrocytes, activated by neuroinflammatory microglia via secretion of Il-1α, TNFα and C1q (5). A1 astrocytes are considered neurotoxic losing their ability to promote neuronal survival and synaptogenesis and exhibit impaired phagocytosis and glutamate uptake (5). Phagocytosis is one of the fundamental elements of the innate immune response. Although microglia are considered the primary phagocytic cell in the CNS, there is increasing appreciation of astrocytes’ role in particular in the phagocytosis of senescent synapses (10–12). Glutamate is the major excitatory neurotransmitter in the CNS. Clearance of glutamate from the synaptic cleft is ensured by a high affinity glutamate uptake system (13,14). Failure to perform this function causes excess glutamate and excitotoxicity (15,16) which is involved in the pathogenesis of several neurological disorders including AD (17). Within the CNS, astrocytes are primarily responsible for glutamate uptake via two glia-specific transporters — excitatory amino acid transporter 1 and 2 (EAAT1 and EAAT2). Ablation of astrocytes in murine models leads to a failure to remove glutamate (18).

Given that ApoE is primarily expressed in astrocytes within the CNS we hypothesised that key physiological astrocytic functions including phagocytosis, glutamate uptake and inflammatory response would be compromised by the *APOE44* genotype and that the A1 phenotype would be potentiated in *APOE44* astrocytes. We created isogenic *APOE* lines using CRISPR Cas-9 gene-editing technology and from them differentiated quiescent astrocytes using a modified protocol. Astrocytes were then stimulated to induce an astrogliotic, A1 phenotype simulating the process of CNS insult, microglial activation and microglial-astrocyte cross-talk (5).

We confirmed that *APOE44* astrocytes have distinct phenotypes in key homeostatic functions both at rest and in their A1 state and that these may be linked by a common NF-κB pathway.

## 2. Materials and Methods

### 2.1. Generating Isogenic APOE4 iPSCs

CRISPR/Cas9 gene editing was used to generate *APOE34* and *APOE44* isogenic lines from human embryonic stem cells (hESC) with an *APOE33* genotype. Editing at the APOE the rs429358 SNP locus was introduced by using cellular homology-directed repair (HDR). A single gRNA sequence (5′-GCGGACATGGAGGACGTGtG**CGG**-3′) and a 98-bp repair template containing the desired edit: (5′-GCGGGCACGGCTGTCCAAGGAGCTGCAGGCGGCGCAGGCCCGGCTGGGCGCGGA<u>T</u>A TGGA<u>A</u>GACGTG<u>c</u>G**<u>T</u>GG**CCGCCTGGTGCAGTACCGCGGCGAGG-3’) provided a template for HDR. The HDR template contained three silent mutations (underlined), one of which was in the PAM sequence (shown in bold) to prevent further excision, and introduced sequences that were utilised during screening.

crRNA and tracrRNA-ATTO-550 (IDT) were combined in nuclease-free duplex buffer (IDT), annealed (95 °C, 2Lminutes), combined with Cas9 (IDT) and incubated (room temperature, 20Lminutes) to form a ribonucleoprotein (RNP). hESCs were nucleofected with RNPs using the 4D-Nucleofector and P3 Primary Cell 4D-Nucleofector X Kit and program CB150 (Lonza). After 24□hours, iPSCs were sorted on the FACSAria Fusion to obtain ATTO-550 positive cells, which were plated as single cells. After 7□days, individual colonies were manually dislodged and plated into single wells of a 96-well plate, which, after 7□days, were passaged into replicate plates using Gentle Cell Dissociation Reagent (STEMCELL Technologies). For screening, DNA was extracted and PCR amplified using a primer pair created to recognise when the HDR template had been incorporated by designing a reverse primer lying over the edit site including three silent mutations and the intended mutation. This created a 193bp product which was visualized on a 1.5% agarose gel on the Gel Doc XR system (Bio-Rad). DNA from clones that were positive for edited product underwent a further PCR using primers flanking the edit site to create a 250bp region of exon 4 of *APOE*; 5′-ACAAATCGGAACTGGAGGAACAA -3′ and 5′-TTCTGCAGGTCATCGGCAT -3′. Sanger sequencing was used to confirm successful editing.

### 2.2. Differentiation of astrocytes

Neural progenitor cells (NPCs) were differentiated using well described protocols (100). NPCs were initially plated on poly-d-lysine laminin coated plates with Sciencell Astrocyte Medium (Cat: 1801). Subsequent passages were onto Geltrex (Fisher Scientific) coated plates with Sciencell Astrocyte Medium. After three weeks, the medium was transitioned to one based on a publication by Serio and colleagues (101) containing 1% FBS and the growth factors BMP4 and CNTF each at a concentration of 20ng/ml; this was continued for a further 5 weeks. Fourth and subsequent passages were onto uncoated plates. Together with robust trituration during passaging, this ensured elimination of neuronal cells. At 12 weeks, growth factors were withdrawn and astrocytes were gradually transitioned to a serum free medium until experimentation.

### 2.3. Induction of astrogliosis

Induction of astrogliosis was undertaken using the combination of cytokines set out in Liddelow and colleagues’ 2017 publication (5). Cells were first plated at the appropriate density 3-7 days prior to the particular experiment in serum free media. 24 hours prior to the experiment, media was aspirated from the wells and cells were washed twice with warmed DPBS. Media containing Il-1α (3ng/ml; Sigma-Aldrich, I13901), TNFα (30ng/ml; NEB 8902SF) and C1q (400ng/ml; MyBioSource, MBS143105) was then added to each well. After 24 hours, media was aspirated, cells washed twice with DPBS and harvested as appropriate.

### 2.4. Immunocytochemistry

Cells were fixed with 4% paraformaldehyde in PBS plus 15 min at room temperature (RT), and washed three times with PBS. Cells were blocked in 10% donkey or goat serum with 0.1% Triton X-100 in PBS for 1h at RT, and subsequently incubated with primary antibody in 3% donkey or goat serum with 0.1% Triton X-100 in PBS overnight at 4°C. Primary antibodies are detailed in Table S1. Cells were washed three times with PBS and incubated with the respective secondary antibody at 1:500 in 3% donkey serum with 0.1% Triton X-100 in PBS for 2h at RT (anti-rabbit Alexa Fluor 568 and anti-mouse Alexa 488, both from donkey, Jackson Immunoresearch). Cells were washed once and incubated with DAPI (40,6-diamidino-2-phe-nylindole, Sigma-Aldrich; 1/3000) for 5 min at RT. Cells were washed twice with PBS before mounting with Fluorescent Mounting medium (Dako).

Stained cells were imaged using a Leica DM6000B inverted microscope. For quantification, random fields were acquired for each well at a 20x magnification. Cell counting of DAPI and most nuclear markers was performed using the ITCN (Image-based tool for counting nuclei) Plugin for ImageJ developed by Thomas Kuo and Jiyun Byun. Images were converted to eight-bit greyscale and inverted before using ITCN. Cell detection was performed by detecting dark peaks with parameters optimised to ensure accurate cell capture.

### 2.5. qPCR

Total RNA was isolated using the Invitrogen PureLink RNA Mini Kit (cat. no. 12183018A) according to the manufacturer’s instructions. RNA concentration and purity were determined using a Nanodrop spectrophotometer. cDNA was prepared from 0.75□μg of RNA. RNA was reverse-transcribed using The High Capacity cDNA Reverse Transcription Kit (Applied Biosystems Cat no. 4368814) according to the manufacturer’s instructions.

Primers for quantitative PCR were designed to be intron-spanning in order to avoid amplification of DNA (see Supplementary Table S1). The reaction was undertaken using SyGreen Blue Mix Hi Rox kit (PCRBiosystems Cat no PB20.15-20). Each sample was run in triplicate. For each reaction, 10 μl of 2X qPCR SyGreen Blue Mix, 0.8μl of both primers (from 10μM stock) for the gene of interest and 75ng of cDNA were combined and made up to 20μl with ddH2O. The qPCR reaction was run on a Bio-Rad CFX Connect Real-Time System. The program used included an initial incubation at 95°C for 2 mins, followed by 40 cycles of 95°C for 30 secs and 60°C for 15 secs. For every primer, a melt curve was generated to check for the specificity of the product; any primers that generated more than one peak were discarded.

### 2.6. Protein measurement

Human ApoE protein was quantified using the Invitrogen Human ApoE ELISA following the manufacturers guidelines (Invitrogen. Cat no. EHAPOE).

Other proteins were quantified using standard immunoblotting techniques. Cultured cells were washed twice in DPBS and lysed on ice using RIPA buffer (Abcam) supplemented with a protease and phosphatase inhibitors cocktail (Sigma). Cell lysates were centrifuged for 15 minutes at 12,000g and the resulting supernatant combined with 1X Bolt®LDS Sample Buffer (Thermo Fisher) and 1X Bolt®Sample Reducing Agent (Thermo Fisher) and denatured at 97°C for 5 minutes.

Next, 20 μg of cell protein extract per sample was separated on 4-12% Bolt® Bis-Tris Plus gels (ThermoFisher) using MOPS SDS NuPAGE Running Buffer (Invitrogen) at 150V for 120□minutes then transferred to PVDF membranes (Sigma-Aldrich) using NuPAGE Transfer Buffer (Invitrogen) at 120□V for 105Lminutes. The membrane was blocked in Protein-Free (TBS) Blocking Buffer (ThermoFisher) and incubated overnight at 4°C with primary antibodies (see Supplementary Table S2).

Antibodies were detected with fluorophore conjugates IRDye 800CW donkey anti-guinea pig IgG secondary antibody, IRDye 800CW goat anti-rabbit IgG secondary antibody IRDye 680RD donkey anti-chicken IgG secondary antibody and 680RD goat anti-mouse IgG secondary antibody (all LI-COR Biosciences, 1:10,000). Immunoblots were visualised using the Odyssey CLx Imaging System using GAPDH as a loading control. The integrated density for bands from one or two replicates each from three independent experiments (n=5 or 6) was measured. Unpaired, two-tailed t-tests were performed to compare *APOE33* and *APOE44* samples.

### 2.7. Cytokine array

A Human Cytokine Antibody Array (Abcam #ab133997) was used according to manufacturer’s instructions. Astrocytes were transitioned to serum free medium at least one week prior to A1 induction; 2mls of treated (as per section 2.3) or untreated medium was added to each well of a 12-well plate. After 24 hours, supernatant was transferred to fresh Eppendorf tubes, and cells were harvested for protein samples. Supernatant was centrifuged at 12,000g for 5 min at 4°C and kept at -80°C until experimentation. Supernatant from 3 independent experiments and 2 biological replicates per experiment was used (n=6 per condition) with 3 separate technical replicates (wells) pooled for each biological sample. Densitometry analysis was calculated using Image J; each spot is duplicated on the test membrane so results represent the mean of these two values. Results were normalised to the array’s internal protein control.

### 2.8. Lipid extraction and oxylipin measurement

The same supernatants used for the cytokine array were also used for oxylipin analysis. Lipid extraction and oxylipin measurements were carried out in the laboratory of Professor Val O’Donnell as previously described (102,103). Samples were spiked with 2.1-2.9ng of PGE_2_-d4, PGD_2_-d4, 20-HETE-d6, 5-HETE-d8, 12-HETE-d8, 15-HETE-d8, 13-HODE-d4 and TXB_2_-d4 standards (Cayman Chemical) prior to extraction. Oxylipins were extracted by adding a 2.5 ml solvent mixture (1 M acetic acid/isopropanol/hexane; 2:20:30, v/v/v) to 1 ml supernatants in a glass extraction vial and vortexed for 30 sec. 2.5ml hexane was added to samples and after vortexing for 30 seconds, tubes were centrifuged (500 g for 5 min at 4 ***°***C) to recover lipids in the upper hexane layer (aqueous phase), which was transferred to a clean tube. Aqueous samples were re-extracted as above by addition of 2.5 ml hexane, and upper layers were combined. Lipid extraction from the lower aqueous layer was then completed according to the Bligh and Dyer technique. Specifically, 3.75ml of a 2:1 ratio of methanol:chloroform was added followed by vortexing for 30 secs. Subsequent additions of 1.25ml chloroform and 1.25ml water were followed with a vortexing step for 30 seconds, and the lower layer was recovered following centrifugation as above and combined with the upper layers from the first stage of extraction. Solvent was dried under vacuum and lipid extract was reconstituted in 100μl HPLC grade methanol. Lipids were separated by liquid chromatography (LC) using a gradient of 30-100% B over 20 minutes (A: Water:Mob B 95:5 + 0.1% Acetic Acid, B: Acetonitrile: Methanol – 80:15 + 0.1% Acetic Acid) on an Eclipse Plus C18 Column (Agilent), and analysed on a Sciex QTRAP^®^ 6500 LC-MS/MS system. Source conditions: TEM 475°C, IS -4500, GS1 60, GS2 60, CUR 35. Lipid were detecting using MRM monitoring with the parent to daughter ion transitions detailed in Supplementary Table S4. Deuterated internal standards were monitored using parent to daughter ions transitions of: TXB2-d4 [M-H]-373.2/173.1, PGE2-d4 and PGD2-d4 [M-H]-355.2/275.1, 5-HETE-d8 [M-H]-327.2/116.1, 12-HETE-d8 [M-H]-327.2/184.1, 15-HETE-d8 [M-H]-327.2/226.1, 13-HODE-d4 [M-H]-299.2/198.1 and 20-HETE-d6 [M-H]-325.2/281.1. Chromatographic peaks were integrated using Multiquant 3.0.2 software (Sciex). The criteria for assigning a peak was signal:noise of at least 5:1 and with at least 7 points across a peak. The ratio of analyte peak areas to internal standard was determined and lipids quantified using a standard curve generated and run at the same time as the samples. All oxylipins were detected in negative ion mode as [M-H]-ions.

### 2.9. Phagocytosis assay

Assessment of phagocytosis was undertaken with the Vybrant Phagocytosis Assay kit (Thermofisher cat: V6694) according to manufacturer’s instructions. Astrocytes were plated in 96-well plates at a density of 30×10^3^/cm^2^ 48 hours prior to testing. As a negative control, wild-type (*APOE33*) astrocytes were treated with the cell-cycle inhibitor Cytochalasin D (5ug/ml for 30 min at 37°C; Thermofisher) which inhibits phagocytosis.

Cells were incubated for 2 hours with 100 μl of fluorescein-labelled E. coli Bioparticles. The E. coli suspension was aspirated, and 100 μl of Trypan Blue was added for 1 minute at RT to quench extracellular fluorescence. Following Trypan Blue aspiration, fluorescence was measured (480 nm excitation/520 nm emission) using a microplate reader.

Average fluorescence intensity values from groups of 3 replicates of negative control, positive control and experimental samples per differentiation (n=9) were used. The Net Experimental Reading was obtained by subtracting the average fluorescence intensity of the negative-control wells (Cytochalasin D treated) from the experimental wells. Results were then normalised to protein levels assessed using a BCA assay. The phagocytosis response was expressed as follows:

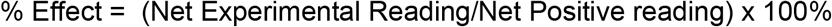

Where the quiescent *APOE33* sample was the Net Positive Reading. Data were analysed with the two-way paired or unpaired student’s t-test as appropriate.

### 2.10. Glutamate uptake assay

Differentiated astrocytes were plated at a density of 50×10^3^/cm^2^ onto Geltrex coated 48-well plates in serum free media 3 days prior to experimentation. Cells were washed with HEPES buffer (Thermofisher) and then starved in HBSS media without calcium and magnesium (Thermofisher) for 30 minutes. After 30 minutes, the HBSS was aspirated and replaced with HBSS with calcium and magnesium plus glutamate (50μM: Sigma) which was also added to cell-free wells to determine the starting glutamate concentration. After 60 minutes at 37°C, samples were aspirated from each well and kept on ice for analysis with a colorimetric glutamate assay kit (Sigma-Aldrich; MAK004), according to the manufacturer’s instructions. Samples of HBSS without glutamate were run as negative controls and subtracted from all readings to obtain a corrected measurement for each sample. Final concentrations for each sample were subtracted from initial concentrations and percentage uptake calculated. N=6 per genotype per condition (2 differentiations, 3 biological replicates for each) with each sample was measured in duplicate. Data were analysed with a two-way paired or unpaired t-test as appropriate.

### 2.11. Statistics

All statistical analysis was carried out using GraphPad PRISM or SPSS versions 26-28. All data are expressed as mean ± standard deviation. Data were analysed with the two-way paired t-test (for quiescent versus A1) or unpaired t-test (for *APOE33* versus *APOE44*). For screening assays such as the cytokine and oxylipin arrays, statistical significance was determined using an error probability level of p <0.05 corrected by a false discovery rate (FDR) analysis (Benjamini Hochberg method). The statistical test used, value of n, and meaning of n are indicated in the figure legends and the corresponding methods section. Statistical significance is defined as p<0.05 (* = p<0.05 ; ** = p<0.01, *** = p<0.001, **** = p<0.0001).

## 3. Results

### 3.1. Creation and verification of isogenic *APOE* astrocytes

CRISPR/Cas9 gene editing was used to generate *APOE34* and *APOE44* human embryonic stem cells (hESCs) from parental hESC7 APOE33 cells using the guide RNA and repair oligonucleotide sequences shown (Figure 1 A). Successful editing resulted in a thymine to cytosine change at the rs429358 locus and a corresponding Cys112Arg substitution in the *APOE* protein product, which was verified by Sanger sequencing of colonies derived from single edited cells (Figure 1 B). Whole-exome sequencing of edited line and parental iPSC lines showed no unintended off-target coding mutations, while karyotyping analysis revealed no chromosomal changes in the isogenic lines. Both parental and genome-edited hESCs maintained comparable expression of pluripotency markers (Figure 1 C).

**Figure 1:**
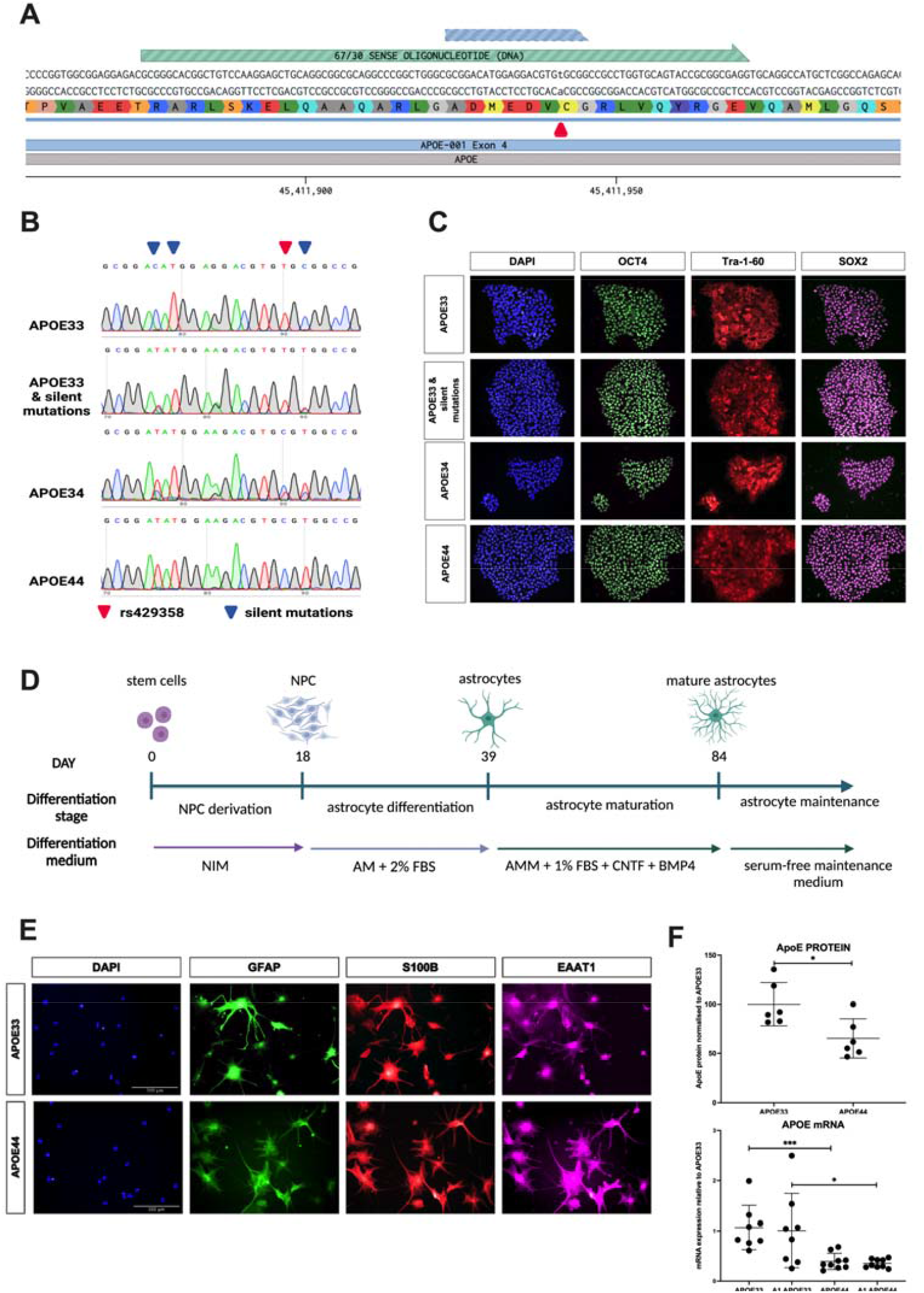
creation and validation of isogenic lines. A) Guide RNA (blue) & repair oligonucleotide (green) sequences for editing the rs429358 locus (red arrow). B) Sanger sequencing of isogenic lines showing successful editing at the rs429358 locus resulting in a thymine to cytosine change and a corresponding Cys112Arg substitution in the *APOE* protein product. C) Immunocytochemistry demonstrating comparable expression of the pluripotency markers OCT4, Tra-1-60 and SOX2 in parental and genome-edited hESCs. D) Schematic for differentiation of quiescent astrocytes: stem cells to neuronal precursor cells (NPCs) then onto mature astrocytes. E) Astrocytes fixed and stained with astrocytic markers GFAP, EAAT1 and S100B at 18 weeks. F) Protein and mRNA expression of astrocytes. Protein was measured using ELISA (n=6, 2 separate differentiations, 3 technical replicates). mRNA using qPCR analyses; n=9, 3 separate differentiations, 3 technical replicates (each technical replicate in triplicate).

Quiescent astrocytes were differentiated as outlined in the methods section, first to neuronal precursor cells (NPCs) then onto mature astrocytes (Figure 1 D). All wild-type and isogenic lines showed expression of the NPC markers nestin, PAX6, OTX2 and FOXG1. Quantification of cells expressing the nuclear stains PAX6, FOXG1 and OTX2 as a percentage of the total expressing DAPi was undertaken to ensure that progression to the NPC stage was comparable in WT and isogenic lines. NPCs were then differentiated until astrocytes were sufficiently mature to survive transition to a serum-free, basal maintenance medium (approximately 12 weeks). Astrocytes were then kept in serum-free maintenance medium until experimentation. Astrocytes were fixed for immunocytochemistry and stained with astrocytic antibodies GFAP, EAAT1 and S100B (Figure 1 E). qPCR of the same astrocytic markers was also undertaken with the addition of the neuronal marker MAP2 which was not detected in our cell cultures.

Isogenic cell lines were validated by measuring the gene and protein expression of *APOE*. In keeping with previous studies, ApoE protein and *APOE* mRNA were significantly reduced in *APOE44* astrocytes, indicating that the APOE4 variant can negatively impact its own transcription, as previously found (19) (Figure 1 F).

### 3.2. Inflammatory profile

A1 astrocytes were induced using the methods described by Liddelow et al (5); briefly, media containing IL-1α (3ng/ml), TNFα (30ng/ml) and C1q (400ng/ml) was added 24 hours prior to experimentation. Cell lysates and conditioned media were collected after 24 hours. The key marker of reactive astrocytes is the complement component C3 which is specifically upregulated in A1 astrocytes but not in quiescent or A2 reactive astrocytes (5). Gene expression of *C3* (Figure 2 A) was measured to ensure that the induction process was effective. As anticipated, *C3* expression was significantly higher in A1 astrocytes than their quiescent counterparts for both *APOE33* (p=<0.0001) and *APOE44* cells (p=<0.0001). Expression of *C3* was also significantly higher in A1 *APOE33* astrocytes compared to A1 *APOE44* astrocytes (*APOE33* 247.7, *APOE44* 123.1; p=0.0011). CD44 another marker of astrocyte activation (20) was also found to be significantly increased in A1 *APOE33* cells compared to their quiescent counterparts (*APOE33* 1.036, A1 *APOE33* 1.794; p=0.0129 but no such difference was observed in *APOE44* cells (*APOE44* 1.141, A1 *APOE44* 1.344; p=0.3220) however this may be due to the variability of the results for this group.

**Figure 2:**
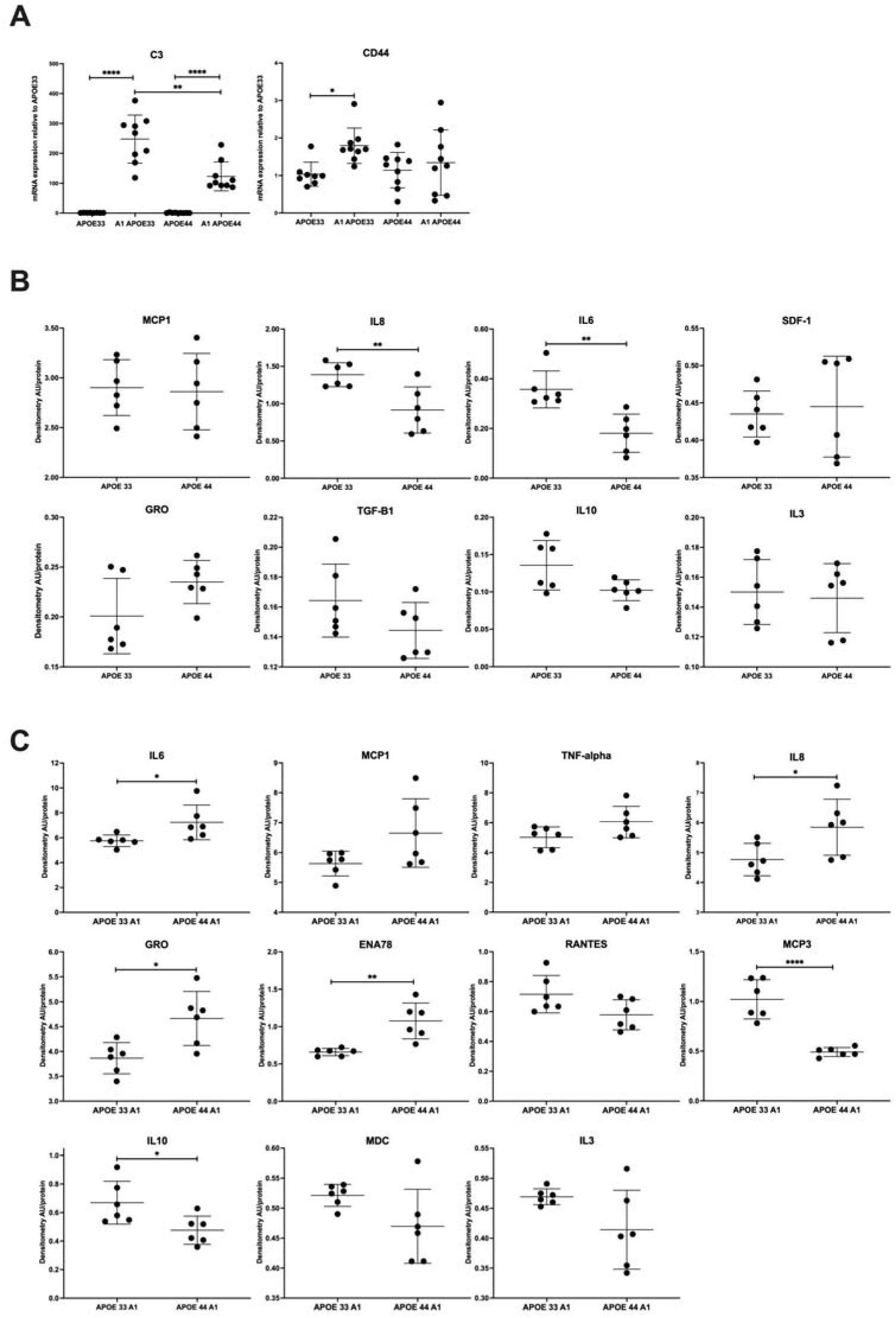
C3, CD44 and cytokine/chemokine expression. All data are presented as mean ± SD using paired or unpaired two-tailed Student’s t test. AU = arbitrary units. A) Quantification of C3 and CD44 mRNA levels using qPCR analyses. n=9, 3 differentiations, 3 technical replicates (each technical replicate in triplicate). B&C) Quantification of cytokines/chemokines in the supernatants of untreated & A1 *APOE33* and *APOE44* astrocytes normalised to protein control (n=6, 3 independent differentiations, 2 technical replicates for each). Cytokines are presented in order of abundance. All cytokines detected are shown for untreated samples, those with an uncorrected p-value <0.1 are shown for A1 samples. Full results may be found in Supplementary data.

When astrocytes undergo astrogliosis, proliferation and release of factors such as cytokines, chemokines (chemotactic cytokines) and growth factors is enhanced (21). The secretory profile of quiescent and A1 astrocytes was examined using cytokine and oxylipin arrays; the full list of oxylipins tested is in Table S4. Cytokines and chemokines were measured using a cytokine array (Abcam) according to the manufacturer’s instructions. It showed that 8 cytokines/chemokines were present in quiescent samples (Figure 2 B). In order of intensity these were: MCP1, IL8, IL6, SDF-1, IL3 GRO (CXCL1), TGFB1, IL10 and IL3. There were significant differences in IL6 (*APOE33* 0.3571, *APOE44* 0.1805; p=0.0024), and IL8 (*APOE33* 1.3890, *APOE44* 0.9155; p=0.0077), which were, unexpectedly, lower in *APOE44* astrocytes; both remained significant after False Discovery Rate (FDR) correction for multiple testing. Differences in IL10 and GRO were approaching significance after FDR.

In A1 astrocytes, a further 8 chemokines/cytokines were expressed yielding a total of 16; these are shown in order of intensity (Figure 2 C). Two targets were significantly increased in *APOE44* astrocytes after False Discovery Rate (FDR) correction for multiple testing: ENA (*APOE33* 0.659, *APOE44* 1.077; p=0.0019) and GRO (*APOE33* 3.864, *APOE44* 4.664; p=0.011). IL6 (*APOE33* 5.759, *APOE44* 7.231; p=0.0336) and IL8 (*APOE33* 4.767, *APOE44* 5.847; p=0.0345) did not remain significant after FDR correction. One target was significantly decreased in *APOE44* astrocytes MCP3 (*APOE33* 1.0210, *APOE44* 0.4918 p=<0.0001); IL10 (*APOE33* 0.6695, *APOE44* 0.4761; p=0.0249) did not remain significant after FDR correction. Five other targets were approaching significance: two showing increases in *APOE44* astrocytes (MCP1 and TNF-alpha) and three showing a decrease (RANTES, MDC and IL3).

Overall, the pattern of expressed cytokines in our quiescent astrocytes was in agreement with previously published data using human fetal astrocytes (22) and differentiated astrocytes (23–25). In A1 astrocytes, the pattern in *APOE44* astrocytes was broadly pro-inflammatory with increases in the pro-inflammatory cytokines ENA, GRO, IL6 and IL8 and decreases in the anti-inflammatory cytokine IL10. However, the finding of lower levels of IL6 and IL8 in quiescent astrocytes was unexpected and may reflect the efficacy of our method for production of genuinely quiescent astrocytes.

The same samples used for cytokine analysis were also analysed for oxylipin levels 24 hours after A1 activation using liquid chromatography/mass spectrometry (LC/MS). Oxylipins (oxygenated polyunsaturated fatty acids) are bioactive lipid mediators synthesised from polyunsaturated fatty acids (PUFAs) by cyclooxygenase (COX), lipoxygenase (LOX) & cytochrome P450 (CYP) activity (26). These are key regulators of peripheral inflammatory responses but are less characterised in the brain. Nineteen oxylipins were generated by quiescent cells; with 7 showing significant *APOE*-genotype dependent differences after FDR correction (Figure 2 D): 5-HETE (*APOE33* 0.0314ng/ml, *APOE44* 0.0595ng/ml; p=<0.0001); 5-HEPE (*APOE33* 0.0639ng/ml, *APOE44* 0.1049ng/ml; p=0.0002); 14,15-DiHETrE (*APOE33* 0.0051ng/ml, *APOE44* 0.00330ng/ml; p=0.0004); 11-HETE (*APOE33* 0.0399ng/ml, *APOE44* 0.0315ng/ml; p=0.0025); 13-HODE (*APOE33* 1.0350ng/ml, *APOE44* 0.8501ng/ml; p=0.00108); 11-HEPE (*APOE33* 0.0393ng/ml, *APOE44* 0.0315ng/ml; p=0.0159), and 9,10-DiHOME (*APOE33* 0.0231ng/ml, *APOE44* 0.0206ng/ml; p=0.0197). All were lower in *APOE44* astrocytes except 5-HETE and 5-HEPE, which were elevated.

In A1 astrocytes, 23 lipids were synthesised (4 of which were expressed in A1 but not quiescent astrocytes). The pattern of increased and decreased oxylipins after treatment was entirely consistent with published data (27). Twelve lipids showed *APOE*-genotype dependent differences, eleven of which remained significant after FDR correction. Of these, all were lower in *APOE44* cells except PGF2α (Figure 2 D): PGE2 (*APOE33* 0.7927ng/ml, *APOE44* 0.3022ng/ml; p=<0.0001); PGE3 (*APOE33* 0.0241ng/ml, *APOE44* 0.0004ng/ml; p=<0.0001); 13-HDOHE (*APOE33* 0.0357ng/ml, *APOE44* 0.0028ng/ml; p=<0.0001); PGF2α (*APOE33* 0.3522ng/ml, *APOE44* 0.5312ng/ml; p=<0.0001); PGE1 (*APOE33* 0.08063ng/ml, *APOE44* 0.0325ng/ml; p=0.0004); 11-HEPE (*APOE33* 0.0550ng/ml, *APOE44* 0.0381ng/ml; p=0.0004); 20-HDOHE (*APOE33* 0.0372ng/ml, *APOE44* 0.0277ng/ml; p=0.0008); 13-HODE (*APOE33* 1.0950ng/ml, *APOE44* 0.8068ng/ml; p=0.0015); 11-HETE (*APOE33* 0.1044ng/ml, *APOE44* 0.0781ng/ml; p=0.0043); 14,15 DiHETrE (*APOE33* 0.0055ng/ml, *APOE44* 0.00044ng/ml; p=0.0085) and 18-HEPE (*APOE33* 0.1293ng/ml, *APOE44* 0.1028ng/ml; p=0.0110). Specialised pro-resolving mediators (SPM), such as resolvins, were not detected in either quiescent or A1 astrocytes.

The majority of the most highly upregulated cytokines in A1 astrocytes are induced by the Nuclear factor-kappa B (NF-κB) transcription factor (28,29) which is also known to influence cyclooxygenase-2 (COX-2), lipoxygenase (LOX) and cytochrome P450 activity (30–33). Given this, we hypothesised that the *APOE* genotype could be associated with alterations in expression of NF-κB proteins. The NF-κB family of transcription factors consists of five different subunits (c-Rel, p65/RelA, p50, RelB, and p52), which interact to form transcriptionally active homo and heterodimers (34). Inactive NF-κB dimers are sequestered in the cytoplasm bound to the IκB inhibitory proteins including IκBα. Using western blotting (Figures 2 F & G) it was found that protein expression of the p50 and p65 subunits were significantly increased in both quiescent and A1 *APOE44* astrocytes (p50 = quiescent *APOE33* 1.489, quiescent *APOE44* 2.152, p=0.0436; A1 *APOE33* 1.666, A1 *APOE44* 3.596; p=0.0001); (p65 = quiescent *APOE33* 0.825, quiescent *APOE44* 2.428, p=0.0027; A1 *APOE33* 2.751, A1 *APOE44* 3.920; p=0.0182). IκBα was also significantly increased in both quiescent and A1 *APOE44* astrocytes (quiescent *APOE33* 0.343, quiescent *APOE44* 1.074, p=0.002; A1 *APOE33* 0.600, A1 *APOE44* 1.130; p=0.0189). There were no significant differences found in the expression of the p105 sub-unit in quiescent astrocytes but it was significantly increased in A1 astrocytes (A1 *APOE33* 0.600, A1 *APOE44* 1.130; p=0.0189).

TNF-α is one of the best known activators of the NF-kB family of transcription factors so it is unsurprising that the induction of the A1 phenotype results in increased expression of cytokines whose genes are regulated by NF-kB. What is interesting, however, is that these effects appear to be influenced by *APOE* genotype and that genotype-dependent differences in the expression of NF-kB proteins are also apparent in quiescent astrocytes although further experimentation to determine activity, not just expression, of NF-kB are required.

### 3.3. Effect of *APOE* genotype on phagocytosis

Phagocytosis was assessed in quiescent and A1 astrocytes (Figure 3 A). It has been shown previously that 24 hour induction of the A1 phenotype reduces astrocytic phagocytosis (5,20) and that phagocytosis is also reduced in astrocytes with *APOE4* genotype (23,35) (although another study failed to find a difference (36)). To our knowledge, this is the first time that the effect of *APOE* genotype on phagocytosis in A1 astrocytes has been investigated. Induction of the A1 phenotype significantly reduced phagocytosis in both *APOE33* and *APOE44* astrocytes. Furthermore, phagocytosis was significantly lower in both quiescent and A1 *APOE44* astrocytes compared with *APOE33* cells (quiescent *APOE33* 100.00%, quiescent *APOE44* 48.63%, p=0.0003; A1 *APOE33* 55.33%, A1 *APOE44* 20.06%; p=0.0006).

**Figure 3:**
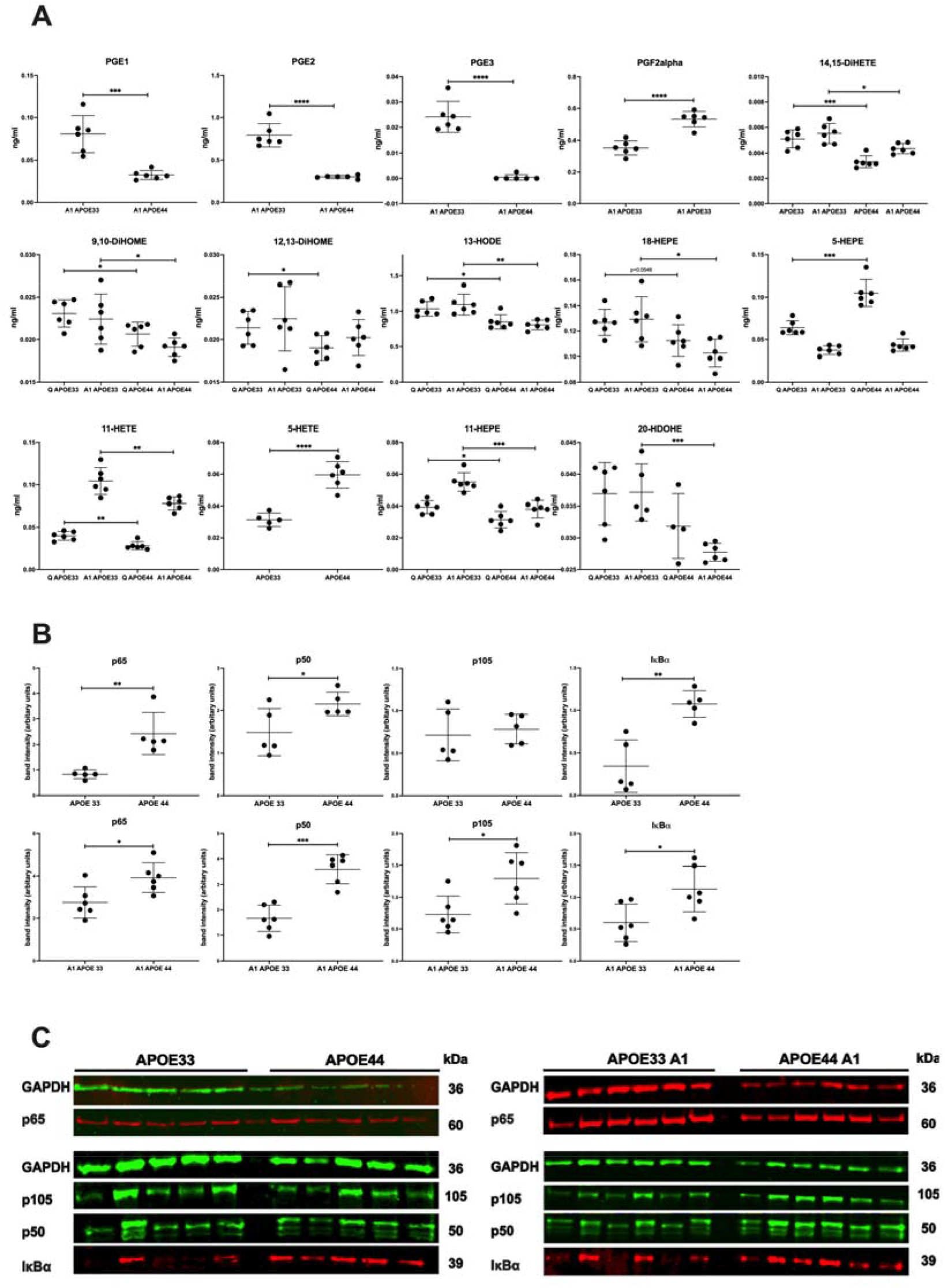
oxylipin synthesis and NF-kB pathway proteins. A) Quantification of oxylipins in the supernatants of untreated and treated (A1) *APOE33* and *APOE44* astrocytes (ng/ml) (n=6, 3 independent differentiations, 2 technical replicates for each). Only those significant after FDR are shown; full results may be found in Supplementary data. B) Quantitative analyses for different proteins in the NF-kB pathway, normalised to GAPDH, in untreated and treated (A1) APOE33 and APOE44 astrocytes (N=5-6, 3 independent differentiations, 1-2 technical replicates). Western blotting was carried out using standard methods. Analysis of means using unpaired two-tailed t-test, error bars ± standard deviation. AU = arbitrary units. C) Western blots for B).

Next, the possible reasons for these differences were investigated. Phagocytic activity of astrocytes is mediated via two main pathways, MEGF10 and MERTK; in a murine model, knockout of either of these phagocytic receptors has been shown to block synaptic pruning (37). Gene expression of *MERTK* and *MEGF10* was calculated in quiescent and A1 astrocytes (Figure 3B). Expression of *MERTK* mRNA was significantly lower in both quiescent and A1 *APOE44* astrocytes compared to their *APOE33* counterparts (quiescent *APOE33* 0.9210, quiescent *APOE44* 0.5630, p=0.0043; A1 *APOE33* 1.0720, A1 *APOE44* 0.6306; p=0.0055). MEGF10 mRNA was significantly lower in A1 *APOE44* astrocytes than in A1 *APOE33* astrocytes but there was no significant difference in quiescent cells (quiescent *APOE33* 1.051, quiescent *APOE44* 0.5882, p=0.0646; A1 *APOE33* 1.1600, A1 *APOE44* 0.4888; p=0.0052).

MERTK works with the integrin pathway to regulate rearrangement of the actin cytoskeleton upon phagocytosis (38). The cytoskeleton of astrocytes is characterised by the expression of three main components: actin, glial fibrillary acidic protein (*GFAP*) and vimentin (*VIM*) (39). GFAP and vimentin are type III intermediate filaments predominantly found in astrocytes. GFAP immunoreactivity is increased in astrogliotic astrocytes which largely reflects changes in the astrocytic cytoskeleton (7). We anticipated, therefore, that expression of intermediate filaments would increase in activated astrocytes to facilitate phagocytosis.

The gene expression of *GFAP* and *VIM* were measured in quiescent and A1 astrocytes; no statistically significant differences were found for *GFAP* but *VIM* mRNA was significantly decreased in quiescent *APOE44* astrocytes, Figure 3B (quiescent *APOE33* 1.062, quiescent *APOE44* 0.6138, p=0.0093; A1 *APOE33* 0.8220, A1 *APOE44* 0.6779; p=0.1218). Western blotting of GFAP protein (Figure 3C&D) showed lower levels in *APOE33* quiescent astrocytes which was approaching significance (*APOE33*= 1.073, *APOE44* 0.831, p=0.0583) but there was significantly increased protein in A1 *APOE33* astrocytes compared to A1 *APOE44* astrocytes (*APOE33*= 1.167, *APOE44* 0.9414, p=0.0253).

### 3.4. Effect of *APOE* genotype on glutamate metabolism

The levels of glutamate uptake in each isogenic line under quiescent and A1 conditions was tested. There were no significant difference between genotypes in quiescent astrocytes but in A1 astrocytes glutamate uptake was significantly lower in *APOE44* cells. Next, the possible mechanisms for the difference in glutamate uptake were investigated. The gene expression of the two main CNS glutamate transporters (*EAAT1* and *EAAT2*) plus glutamine synthetase (*GLUL*) were measured (Figures 4 B-D). There were no differences in the expression of *EAAT2* and *GLUL* but we found *EAAT1* mRNA expression to be significantly reduced in quiescent *APOE44* astrocytes when compared to *APOE33* cells (*APOE33*= 1.012, *APOE44* 0.684, p=0.0049) and approaching significance in A1 astrocytes (*APOE33*= 0.8480, *APOE44* 0.5823, p=0.0805).

**Figure 4:**
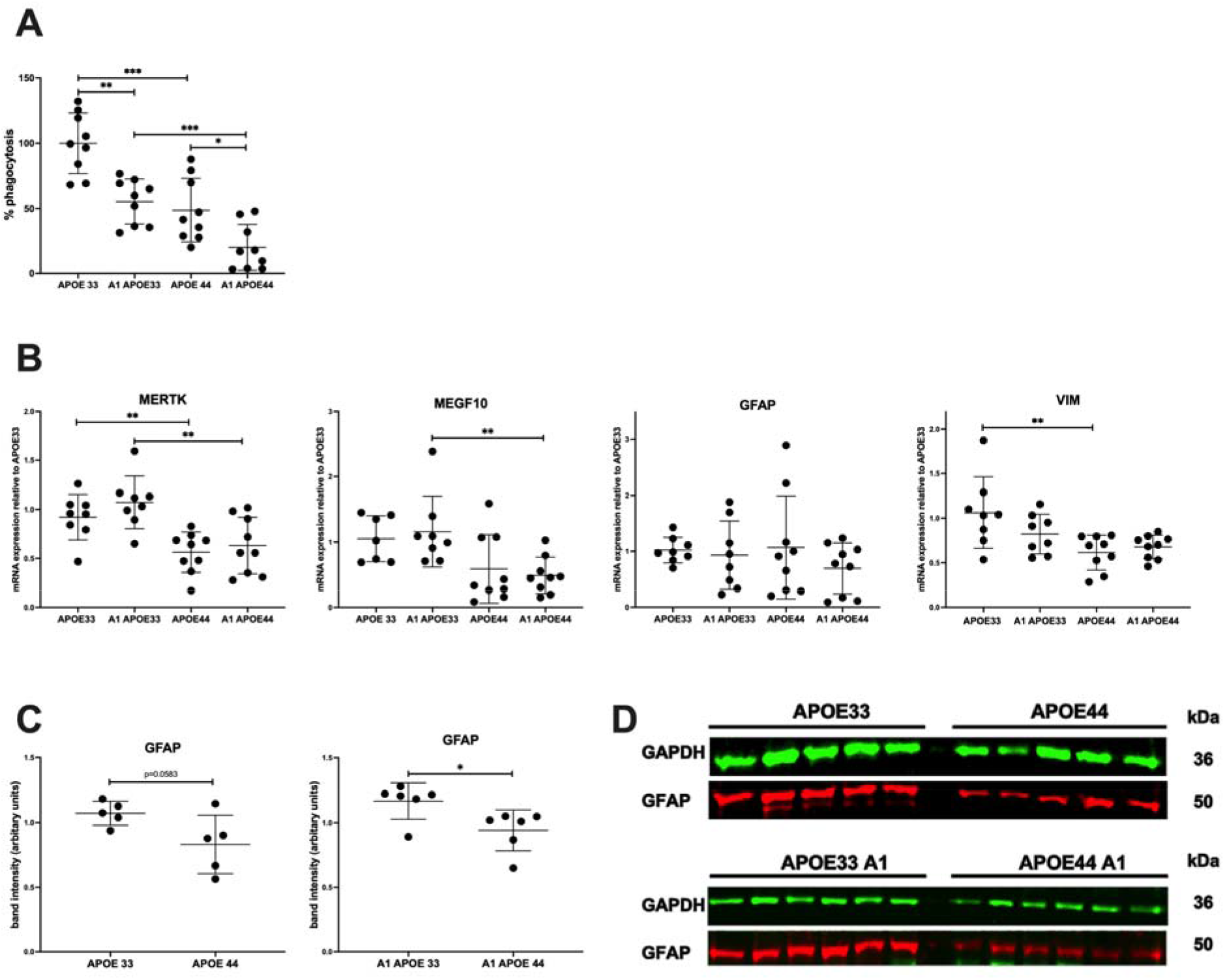
the effect of APOE genotype on phagocytosis. All data are presented as mean ± SD using paired or unpaired two-tailed t-test as appropriate. AU = arbitrary units. A) Quantification of percentage phagocytosis in *APOE33* and *APOE44* astrocytes under quiescent and A1 conditions (N=9, 3 separate differentiations, 3 technical replicates). B) Quantification of *MERTK, MEGF10, GFAP* and *VIM* mRNA levels using qPCR analyses. N=5-6, 3 differentiations, 1-2 technical replicates, each in triplicate. C) Quantitative analyses for GFAP untreated and treated (A1) *APOE33* astrocytes (N=5, 3 differentiations, 1-2 technical replicates) and *APOE44* astrocytes (N=6, 3 differentiations, 2 technical replicates). D) Western blots for C).

**Figure 5:**
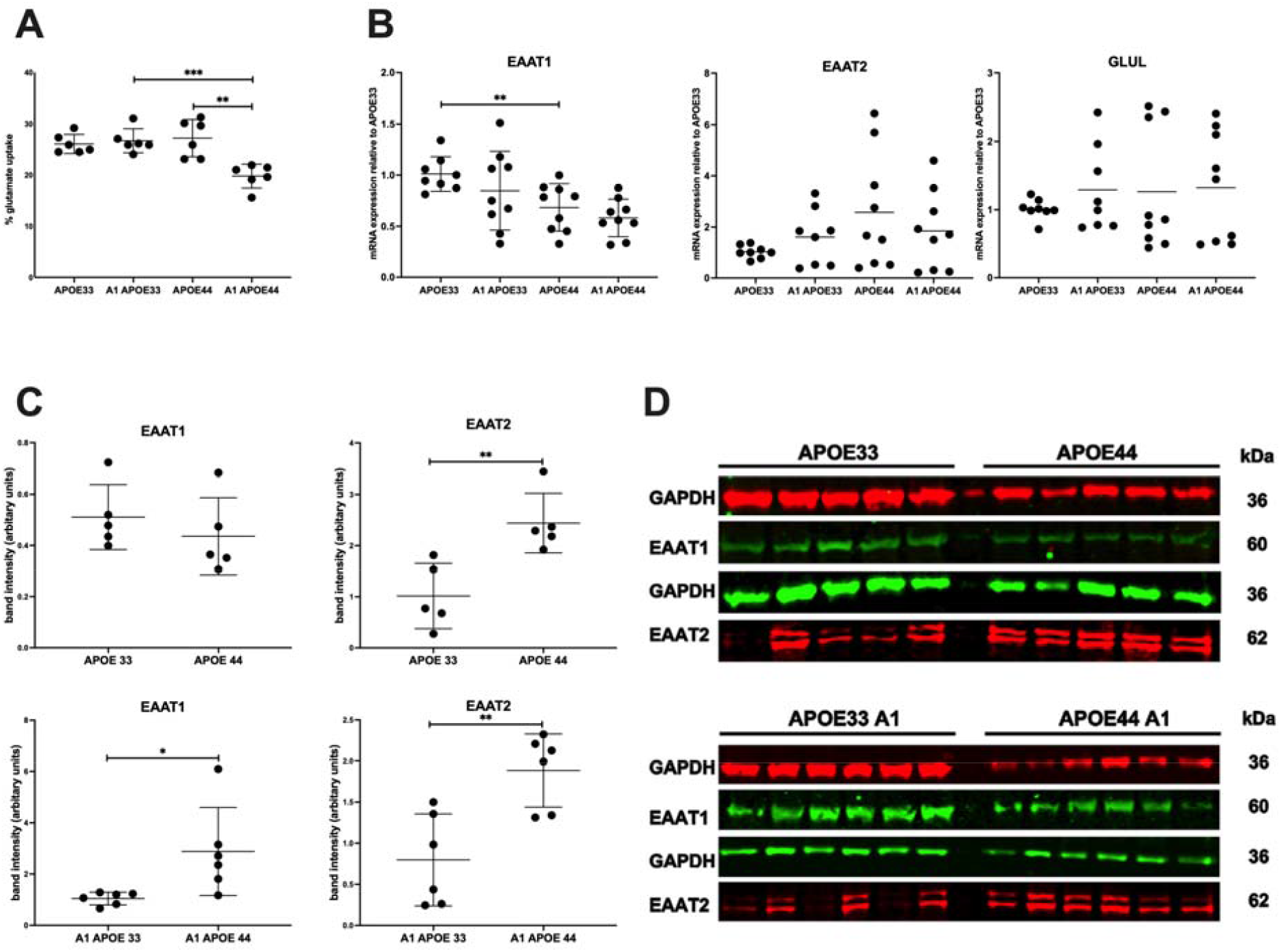
glutamate metabolism. All data are presented as mean ± SD using paired or unpaired two-tailed Student’s t test as appropriate. AU = arbitrary units. A) Percentage glutamate uptake by *APOE* genotypes in quiescent and A1 astrocytes. n=6, 2 differentiations, 3 technical replicates. B) Quantification of EAAT1, EAAT2 and GLUL mRNA levels using qPCR analyses. n=5-6, 3 differentiations, 1-2 technical replicates, each in triplicate. C) Quantitative analyses and western blots for EAAT1 and EAAT2 in quiescent and A1 astrocytes (n=5-6, 3 differentiations, 1-2 technical replicates). D) Western blots for C).

Protein expression of these glutamate receptors was then measured using western blotting. Surprisingly, there was significantly increased expression of the EAAT2 protein in both quiescent *APOE44* astrocytes and A1 *APOE44* astrocytes (quiescent *APOE33* 1.015, quiescent *APOE44* 2.440, p=0.0062; A1 *APOE33* 0.7976, A1 *APOE44* 1.884; p=0.0039). (Figures 4 E & 4 F). For EAAT1, there was no significant difference in quiescent astrocytes but again, unexpectedly, there was significantly increased expression in A1 *APOE44* astrocytes (A1 *APOE33* 1.0460, A1 *APOE44* 2.881; p=0.0270).

## 4. Discussion

Several recent papers using isogenic *APOE* model systems (36,40–44) have implicated *APOE44-*dependent astrocytic phenotypes in the pathogenesis of sporadic Alzheimer’s disease with perturbations of the inflammatory response a common theme. Microglial-astrocyte cross-talk, which stimulates the process of astrogliosis, is a vital part of the inflammatory cascade now considered central to sAD pathogenesis (45).

Chemokine release is part of the process of astrogliosis inducing activation and directional migration of leukocyte subsets to inflammatory sites (46) and facilitating cellular communication between astrocytes, infiltrating leukocytes and other cells thus regulating CNS inflammation (47). The finding of significantly reduced levels of the pro-inflammatory cytokines IL6 and IL8 in untreated astrocytes was unexpected but may indicate that our unstimulated astrocytes were truly quiescent; our protocol was specifically designed to enhance the resting phenotype by complete withdrawal of FBS and the use of the growth factors CNTF and BMP4 (48) highlighting the importance of methods of differentiation.

Furthermore, although IL6 and IL8 are generally considered ‘pro-inflammatory’, at resting concentrations both have neuroprotective functions such as prevention of excitotoxicity, promoting axonal growth, synaptic plasticity and memory consolidation (49,50) which may suggest disruption of the resting *APOE44* phenotype; our oxylipin data are consistent with this. Studies of oxylipin secretion in astrocytes are uncommon (27,51,52) but comparison of published data with our results (Table 1) suggests a clear perturbation in *APOE44* astrocytes; oxylipins that are known to increase in response to lipopolysaccharide (LPS) stimulation (14,15-DiHETrE, 11-HETE, 13-HODE) all have lower basal levels in quiescent *APOE44* astrocytes whereas oxylipins known to decrease in response to LPS stimulation (5-HETE) have significantly higher basal levels in quiescent *APOE44* astrocytes. Interestingly, both 5-HETE and 5-HEPE which had significantly higher basal levels in *APOE44* astrocytes (along with 4-HDOHE which just failed to reach significance after FDR testing) are all products of the enzyme 5-LOX. 5-LOX is the key enzyme involved in the production of pro-inflammatory leukotrienes and also produces reactive oxygen species (ROS) which have long been of interest in sAD (53). 5-LOX levels have been found to be elevated in the brains of AD patients (54,55) and 5-LOX inhibitors are considered potential target for anti-inflammatory drugs (56,57). It has also been shown that p65 interaction with 5-LOX activates NF-κB (33) and that 5-LOX inhibitors ameliorate TNFα-induced cytokine/chemokine release through down-regulation of NF-κB (33,58).

**Table 1:** oxylipin expression in quiescent astrocytes.

| Oxylipin | *APOE44* relative to *APOE33* | Known effect of LPS/TNF- $\alpha$ | p-value | Significant after FDR | Pathway |
| --- | --- | --- | --- | --- | --- |
| 5-HETE | **Increased** | **Decreased** | <0.0001 | Yes | AA/LOX (5-LOX) |
| 5-HEPE | Increased | Unknown | 0.0002 | Yes | EPA/LOX (5-LOX) |
| 14,15-DiHETrE | **Decreased** | **Increased** | 0.0004 | Yes | AA/CYP-EPOXY |
| 11-HETE | **Decreased** | **Increased** | 0.0025 | Yes | AA/LOX (15-LOX) |
| 13-HODE | **Decreased** | **Increased** | 0.0108 | Yes | LA/LOX (15-LOX) |
| 11-HEPE | Decreased | Unknown | 0.0159 | Yes | EPA/LOX |
| 9,10-DiHOME | Decreased | No change | 0.0197 | Yes | LA/CYP-EPOXY |
| 13-HOTrE | Decreased | Unknown | 0.0274 | No | LA/LOX (15-LOX) |
| 4-HDOHE | Increased | Unknown | 0.0314 | No | DHA/LOX (5-LOX) |
| 12,13-DiHOME | Decreased | Unknown | 0.0365 | No | LA/CYP-EPOXY |

**Table 2:** oxylipin expression in A1 astrocytes.

| Oxylipin | *APOE44* relative to *APOE33* | Known effect of LPS/TNF-α | p-value | Significant after FDR | Pathway |
| --- | --- | --- | --- | --- | --- |
| PGE2 | Decreased | Increased | <0.0001 | Yes | AA/COX |
| PGE3 | Decreased | Increased | <0.0001 | Yes | EPA-DHA/COX |
| PGF2α | Increased | Increased | <0.0001 | Yes | AA/COX |
| 13-HDOHE | Decreased | Increased | <0.0001 | Yes | DHA/COX |
| PGE1 | Decreased | Increased | 0.0004 | Yes | LA/COX |
| 11-HEPE | Decreased | Unknown | 0.0004 | Yes | EPA/LOX |
| 20-HDOHE | Decreased | Decreased | 0.0008 | Yes | DHA/CYP |
| 13-HODE | Decreased | Increased | 0.0015 | Yes | LA/LOX |
| 11-HETE | Decreased | Increased | 0.0043 | Yes | AA/COX |
| 14,15-DiHETrE | Decreased | Slight increase | 0.0085 | Yes | AA/CYP-EPOXY |
| 18-HEPE | Decreased | Unknown | 0.011 | Yes | EPA/COX2 or CYP |
| 9-10, DiHOME | Decreased | Decreased | 0.0269 | No | LA/CYP-EPOXY |
| 9-HOTrE | Decreased | Unknown | 0.0414 | No | LA/LOX |

In A1 *APOE44* astrocytes, chemokines showed a pro-inflammatory pattern which accords with previous studies (23,24). The three most upregulated chemokines in *APOE44* astrocytes (GRO (CXCL1), ENA78 and IL8 are known collectively as ‘neutrophil-specific chemokines’ and are primarily expressed by astrocytes in the CNS (59,60). Increased levels of all three chemokines have been found in the CSF and brain parenchyma of patients with AD (61–65). These three chemokines also have a common receptor, CXCR2 (66,67) which has been reported in dystrophic neurites in hippocampal tissue from AD patients (68). It is also upregulated by TNFα and *in vivo* after traumatic brain injury (69). The most differentially expressed chemokine in our samples was monocyte-chemotactic protein 3. MCP3 (CCL7) which was significantly reduced in A1 *APOE44* astrocytes. CCL7 acts as a chemoattractant for several leukocytes, including monocytes but also neutrophils, NK cells and activated T lymphocytes (70). Within the CNS, MCP-3 is produced by astrocytes (71) and is rapidly increased by TNF-α via the NF-κB pathway (72). To our knowledge, decreased expression of MCP-3 has not previously been described in models of Alzheimer’s disease.

Our oxylipin array also showed clear *APOE*-genotype related phenotypes in A1 astrocytes. The top five oxylipins that showed genotype-dependent differences are all formed via cyclooxygenase (COX) activity whose synthesis is mediated by NF-κB but also by pro-inflammatory cytokines (73) suggestive of a vicious cycle of inflammatory activation. Surprisingly, 10 of the 11 significantly different oxylipin levels were lower in *APOE44* astrocytes, the notable exception being PGF2α.

PGE2 and PGF2α (along with 13-HODE) were by far the most abundantly synthesised oxylipins in A1 astrocytes. The *APOE*-dependent differences may reflect variability in the kinetics of oxylipin production between genotypes; while cytokine and chemokine production peaks around 48 to 72 hours post treatment (25), most oxylipin synthesis peaks at approximately 6-12 hours with the oxylipins produced rapidly degraded to other products (74); a time series of oxylipin measurements would confirm this. However, the reversed ratio of PGE2 to PGF2α observed in A1 *APOE44* astrocytes is likely to have pro-inflammatory effects.

While PGE2 is often considered a mediator of active inflammation its effects depend on which of its four main receptors (EP1-4) it acts upon. Importantly it has anti-inflammatory effects including binding to microglial EP2 and EP4 receptors to block LPS induced cytokine synthesis (75) and promotes the induction of suppressive IL-10 (76). PGF2α on the other hand is almost exclusively pro-inflammatory (77). Increased levels of the PGF2α metabolite 13,14-dihydro 15-keto PGF2α has been found in hippocampal pyramidal neurons and reactive astrocytes of Alzheimer disease patients (78). PGF2α is usually rapidly degraded enzymatically with a half-life is less than 1 min in peripheral circulation (79). Its abundance in the *APOE44* samples may reflect perturbation of this key homeostatic mechanism.

There are several pathways by which PGF2α may be synthesised. It begins with the conversion of arachidonic acid through COX activity into the unstable intermediate PGH2 (80). PGH2 can then be converted to prostaglandins (including PGE2) by prostaglandin synthases. PGF2α can be produced from either PGH2 or PGE2 by members of the aldo-keto reductase (AKR) family or PGH2 or PGE2 may interconvert via PGE-F isomerases (81). If PGE2 is reduced, then the higher availability of PGH2 may drive direct conversion of PGF2α; further experimentation will determine which of these pathways may be of relevance.

As noted previously, most cytokines, chemokines and oxylipins produced by astrocytes are targets of the transcription factor NF-κB (22). We found that protein levels of the p50 and p65 subunits and IκBα to be significantly increased in both quiescent and A1 *APOE44* astrocytes which echoes recent findings by Arnaud et al (82). NF-κB is implicated in systemic ageing, inflammation and apoptosis as well as the pathogenesis of several neurodegenerative diseases (28,29). Post-mortem studies of the AD brain generally indicate increased expression and activation of NF-κB particularly in regions preferentially affected in AD such as the hippocampus and entorhinal cortex (83). ApoE4 is known to undergo nuclear translocation binding specifically, and with high affinity, to the promoter regions of 1700 genes many of which have functions implicated in sAD pathogenesis (84). Furthermore, interaction of ApoE4 (but not ApoE3) and the p65 subunit of the NF-κB complex (upregulated in our *APOE44* astrocytes) led to nuclear translocation of p65, suggesting ApoE4 affects NF-κB-mediated gene transcription (85).

*APOE*-dependent differences in NF-κB expression may also be linked to the phenotypes we observed in phagocytosis and glutamate uptake. Astrocytes are particularly important for the elimination of senescent synapses (12) and dystrophic neurites (86). Appropriate clearance of dead cells by phagocytosis is necessary for the development, maintenance, and regeneration of the CNS (87). Precisely how astrocytes contact and take up synapses is still a matter of conjecture (87) but we speculated that expression of intermediate filaments would increase in activated astrocytes to facilitate phagocytosis. While the significantly lower expression of GFAP protein in A1 *APOE44* astrocytes supports this, the prominent reduction of phagocytosis in quiescent *APOE44* astrocytes may be due to other mechanisms as no differences in intermediate filament mRNA or protein were observed (although lower levels of GFAP protein in APOE44 astrocytes was approaching significance). A study of astrocytes in the mediobasal hypothalamus found that chronic upregulation of IKKb/NF-kB (the phenotype we found in our *APOE44* astrocytes) was associated with decreased astrocytic plasticity and shortening of high-order processes. As well as potential impacts on phagocytosis, this phenotype may have implications for synaptic activity as without the support of astrocytic processes, sprouting dendritic filopodia have a short lifespan and tend to retract (88).

While there were no significant *APOE*-dependent differences in glutamate uptake in quiescent astrocytes, glutamate uptake was significantly impaired in *APOE44* A1 astrocytes. Glutamate uptake is undertaken almost exclusively by astrocytes (13,14). EAAT2 performs 90% of glutamate uptake in the mammalian CNS (89) and while it is found throughout the brain, it predominates across the forebrain particularly in the hippocampus (90). Various neurodegenerative diseases have been associated with reduced EAAT2 expression and function including AD (91,92) so the increased expression of EAAT2 may represent a phenotype observed early in the disease process as a similar finding was noted in a murine model of Parkinson’s disease on exposure to alpha-synuclein (93). It is suggested that EAATs may be upregulated to attenuate excitotoxic effects, but over time with disease this upregulation is not sufficient, leading to neuronal loss and eventual reductions in EAAT expression (17). The baseline increase in EAAT2 expression in *APOE44* astrocytes may be linked to increased NF-κB expression as several studies have shown that transcriptional activation of EAAT2 is via NF-κB (94–98) with specific interaction of NF-κB sub-units (notably p65 (97)) with the EAAT2 promoter (96).

In summary, we demonstrate several cellular and transcriptional phenotypes in *APOE44* astrocytes pertinent to the early stages of disease pathogenesis and which may be connected by aberrant NF-kb signaling. This study suggests that therapeutic approaches targeted at restoring NF-kb and inflammatory homeostasis may be beneficial in sAD, particularly in those carrying *APOE4* alleles.

## 5. Conclusion

Expression of inflammatory mediators is vital for regulation of CNS activities, in particular, when the brain is subject to stress. Production of these mediators is under strict control to prevent inflammatory tissue damage. We speculate that this process is compromised in *APOE44* astrocytes leading to perturbations in the finely balanced feedback and synergistic mechanisms that control the inflammatory cascade.

We found that key components of the NF-κB pathway are upregulated in both quiescent and activated *APOE44* astrocytes which is in agreement with recent findings of Arnaud and colleagues (82). Like them we propose that the potentiated NF-κB pathway in *APOE44* astrocytes is central to the derangement of inflammatory phenotypes in both quiescent and A1 astrocytes, rendering them more susceptible to CNS insult. We have also suggested ways in which NF-κB may be relevant to the other *APOE44* astrocytic phenotypes we observed. Multiple studies have demonstrated that signaling through NF-κB in astrocytes contributes to pro-inflammatory responses following injury and that inhibition of NF-κB in astrocytes can promote functional recovery (99). Further exploration of this may provide a fruitful therapeutic target for those possessing *APOE4* alleles.

## Acknowledgments

The authors would like to dedicate this study to Professor Lesley Jones who provided invaluable advice to APR at inception and throughout the project. We also thank Professor Luke O’Neill at Trinity College Dublin for helpful insights into the oxylipin results and members of the Hadyn Ellis Core team for ongoing support. This work was funded by APR’s Alzheimer’s Research UK (ARUK) Clinical Research Fellowship (ARUK-CRF 2016A-1) and the Welsh Clinical Academic Track (WCAT) Programme and supported by the MRC (515781) and the UK Dementia Research Institute (grants 517587 and 5200449) which receives its funding from UK DRI Ltd, funded by the UK Medical Research Council, Alzheimer’s Society and Alzheimer’s Research UK.

## References

1. Husain MA, Laurent B, Plourde M. APOE and Alzheimer’s Disease: From Lipid Transport to Physiopathology and Therapeutics. Front Neurosci. 2021;15(February):1–15.

2. Genin E, Hannequin D, Wallon D, Sleegers K, Hiltunen M, Combarros O, et al. APOE and Alzheimer disease: a major gene with semi-dominant inheritance. Mol Psychiatry. 2011/05/10. 2011 Sep;16(9):903–7.

3. Pitas RE, Boyles JK, Lee SH, Foss D, Mahley RW. Astrocytes synthesize apolipoprotein E and metabolize apolipoprotein E-containing lipoproteins. Biochim Biophys Acta (BBA)/Lipids Lipid Metab. 1987;917(1):148–61.

4. Vasile F, Dossi E, Rouach N. Human astrocytes: structure and functions in the healthy brain. Brain Struct Funct. 2017;222(5):2017–29.

5. Liddelow SA, Guttenplan KA, Clarke LE, Bennett FC, Bohlen CJ, Schirmer L, et al. Neurotoxic reactive astrocytes are induced by activated microglia. Nature. 2017;

6. Jones L, Holmans P a, Hamshere ML, Harold D, Moskvina V, Ivanov D, et al. Genetic evidence implicates the immune system and cholesterol metabolism in the aetiology of Alzheimer’s disease. PLoS One. 2010 Jan;5(11):e13950.

7. Carole Escartin, Elena Galea, András Lakatos, James P. O’Callaghan, Gabor C. Petzold, Alberto Serrano-Pozo, Christian Steinhäuser Andrea Volterra, Giorgio Carmignoto AA, Affiliations. Reactive astrocyte nomenclature, definitions, and future directions. Nat Neurosci. 2021;Feb 15.

8. Sofroniew M V. Astrogliosis. Cold Spring Harb Perspect Biol. 2014;7(2):1–16.

9. Hsu ET, Gangolli M, Su S, Holleran L, Stein TD, Alvarez VE, et al. Astrocytic degeneration in chronic traumatic encephalopathy. Acta Neuropathol. 2018 Dec;136(6):955–72.

10. Chung W-S, Clarke LE, Wang GX, Stafford BK, Sher A, Chakraborty C, et al. Astrocytes mediate synapse elimination through MEGF10 and MERTK pathways. Nature. 2013 Dec;504(7480):394–400.

11. Bellesi M, de Vivo L, Chini M, Gilli F, Tononi G, Cirelli C. Sleep Loss Promotes Astrocytic Phagocytosis and Microglial Activation in Mouse Cerebral Cortex. J Neurosci. 2017 May;37(21):5263–73.

12. Lee JH, Kim J young, Noh S, Lee H, Lee SY, Mun JY, et al. Astrocytes phagocytose adult hippocampal synapses for circuit homeostasis. Nature. 2021;590(7847):612–7.

13. Nicholls D, Attwell D. The release and uptake of excitatory amino acids. Trends Pharmacol Sci. 1990 Nov;11(11):462–8.

14. Lipton SA, Rosenberg PA. Excitatory amino acids as a final common pathway for neurologic disorders. N Engl J Med. 1994 Mar;330(9):613–22.

15. Choi DW. Glutamate neurotoxicity and diseases of the nervous system. Neuron. 1988;1(8):623–34.

16. Doble A. The role of excitotoxicity in neurodegenerative disease: implications for therapy. Pharmacol Ther. 1999 Mar;81(3):163–221.

17. Todd AC, Hardingham GE. The regulation of astrocytic glutamate transporters in health and neurodegenerative diseases. Int J Mol Sci. 2020;21(24):1–32.

18. Delaney CL, Brenner M, Messing A. Conditional ablation of cerebellar astrocytes in postnatal transgenic mice. J Neurosci. 1996 Nov;16(21):6908–18.

19. Lin Y, Seo J, Gao F, Ko T, Yankner BA, Tsai L, et al. APOE4 Causes Widespread Molecular and Cellular Alterations Associated Phenotypes in Human iPSC-Derived Brain Cell Types. Neuron. 2018;1–14.

20. Lilianne Barbar, Tanya Jain, Matthew Zimmer, Ilya Kruglikov, Jessica Sadick, Minghui Wang, Kriti Kalpana, Indigo V.L. Rose, Suzanne R. Burstein, Tomasz Rusielewicz, Madhura Nijsure, Kevin A. Guttenplan, Angelique di Domenico, Gist Croft, Bin Zhang, Hiroko VF. CD49f is a novel marker of functional and reactive human iPSC-derived astrocytes. Neuron. 2020;107(3):436–53.

21. Meeuwsen S, Persoon-Deen C, Bsibsi M, Ravid R, Van Noort JM. Cytokine, chemokine and growth factor gene profiling of cultured human astrocytes after exposure to proinflammatory stimuli. Glia. 2003;43(3):243–53.

22. Choi SS, Lee HJ, Lim I, Satoh JI, Kim SU. Human astrocytes: Secretome profiles of cytokines and chemokines. PLoS One. 2014;9(4).

23. Tcw J, Qian L, Pipalia NH, Chao MJ, Liang SA, Shi Y, et al. Cholesterol and matrisome pathways dysregulated in astrocytes and microglia. Cell. 2022;185(13):2213-2233.e25.

24. de Leeuw SM, Kirschner AWT, Lindner K, Rust R, Budny V, Wolski WE, et al. APOE2, E3, and E4 differentially modulate cellular homeostasis, cholesterol metabolism, and inflammatory response in isogenic iPSC-derived astrocytes. Stem Cell Reports. 2021;17(2021):1–17.

25. Hyvärinen T, Hagman S, Ristola M, Sukki L, Veijula K, Kreutzer J, et al. Co-stimulation with IL-1β and TNF-α induces an inflammatory reactive astrocyte phenotype with neurosupportive characteristics in a human pluripotent stem cell model system. Sci Rep. 2019;9(1):1–15.

26. Funk CD. Prostaglandins and leukotrienes: advances in eicosanoid biology. Science. 2001 Nov;294(5548):1871–5.

27. Chistyakov D V., Gavrish GE, Goriainov S V., Chistyakov V V., Astakhova AA, Azbukina N V., et al. Oxylipin profiles as functional characteristics of acute inflammatory responses in astrocytes pre-treated with IL-4, IL-10, or LPS. Int J Mol Sci. 2020;21(5).

28. Chen CH, Zhou W, Liu S, Deng Y, Cai F, Tone M, et al. Increased NF-κB signalling up-regulates BACE1 expression and its therapeutic potential in Alzheimer’s disease. Int J Neuropsychopharmacol. 2012;15(1):77–90.

29. Bellucci A, Bubacco L, Longhena F, Parrella E, Faustini G, Porrini V, et al. Nuclear Factor-κB Dysregulation and α-Synuclein Pathology: Critical Interplay in the Pathogenesis of Parkinson’s Disease. Front Aging Neurosci. 2020;12(March):1–13.

30. Kaltschmidt B, Linker RA, Deng J, Kaltschmidt C. Cyclooxygenase-2 is a neuronal target gene of NF-kappaB. BMC Mol Biol. 2002 Dec;3:16.

31. Zordoky BNM, El-Kadi AOS. Role of NF-kappaB in the regulation of cytochrome P450 enzymes. Curr Drug Metab. 2009 Feb;10(2):164–78.

32. Navarro-Mabarak C, Mitre-Aguilar IB, Camacho-Carranza R, Arias C, Zentella-Dehesa A, Espinosa-Aguirre JJ. Role of NF-κB in cytochrome P450 epoxygenases downregulation during an inflammatory process in astrocytes. Neurochem Int. 2019;129(June):104499.

33. Jatana M, Giri S, Ansari MA, Elango C, Singh AK, Singh I, et al. Inhibition of NF-κB activation by 5-lipoxygenase inhibitors protects brain against injury in a rat model of focal cerebral ischemia. J Neuroinflammation. 2006;3:1–13.

34. Perkins ND. Achieving transcriptional specificity with NF-κb. Int J Biochem Cell Biol. 1997;29(12):1433–48.

35. Chung W-S, Verghese PB, Chakraborty C, Joung J, Hyman BT, Ulrich JD, et al. Novel allele-dependent role for APOE in controlling the rate of synapse pruning by astrocytes. Proc Natl Acad Sci. 2016;113(36):10186–91.

36. de Leeuw S, Kirschner A, Lindner K, Rust R, Budny V, Wolski W, Gavin A-C NR and TC. APOE2, E3, and E4 differentially modulate cellular homeostasis, cholesterol metabolism, and inflammatory response in isogenic iPSC-derived astrocytes. Stem Cell Reports. 2022;17(January 11):110–26.

37. Chung W-S, Clarke LE, Wang GX, Stafford BK, Sher A, Chakraborty C, et al. Astrocytes mediate synapse elimination through MEGF10 and MERTK pathways. Nature. 2013;504(7480):394–400.

38. Jung YJ, Chung WS. Phagocytic roles of glial cells in healthy and diseased brains. Biomol Ther. 2018;26(4):350–7.

39. Chiu FC, Norton WT, Fields KL. The cytoskeleton of primary astrocytes in culture contains actin, glial fibrillary acidic protein, and the fibroblast-type filament protein, vimentin. J Neurochem. 1981 Jul;37(1):147–55.

40. Zhao J, Davis MD, Martens YA, Shinohara M, Graff-Radford NR, Younkin SG, et al. APOE ε4/ε4 diminishes neurotrophic function of human iPSC-derived astrocytes. Hum Mol Genet. 2017;26(14):2690–700.

41. Tcw J, Liang SA, Qian L, Pipalia NH, Chao MJ, Shi Y, et al. Cholesterol and matrisome pathways dysregulated in human APOE □4 glia. 2019;4.

42. Lin YT, Seo J, Gao F, Feldman HM, Wen HL, Penney J, et al. APOE4 Causes Widespread Molecular and Cellular Alterations Associated with Alzheimer’s Disease Phenotypes in Human iPSC-Derived Brain Cell Types. Neuron. 2018;98(6):1141-1154.e7.

43. Wang C, Najm R, Xu Q, Jeong D eun, Walker D, Balestra ME, et al. Gain of toxic apolipoprotein E4 effects in human iPSC-derived neurons is ameliorated by a smallmolecule structure corrector. Nat Med. 2018;24(May):1–11.

44. Blanchard JW, Akay LA, Davila-Velderrain J, von Maydell D, Mathys H, Davidson SM, et al. APOE4 impairs myelination via cholesterol dysregulation in oligodendrocytes. Nature. 2022 Nov;611(7937):769–79.

45. Singh D. Astrocytic and microglial cells as the modulators of neuroinflammation in Alzheimer’s disease. J Neuroinflammation. 2022 Aug;19(1):206.

46. Luster AD. Chemokines--chemotactic cytokines that mediate inflammation. N Engl J Med. 1998 Feb;338(7):436–45.

47. Sternberg EM. Neural-immune interactions in health and E M Sternberg Find the latest versionL: Perspectives Series□: Cytokines and the Brain. 1997;100(11):2641–7.

48. Magistri M, Khoury N, Mazza EMC, Velmeshev D, Lee JK, Bicciato S, et al. A comparative transcriptomic analysis of astrocytes differentiation from human neural progenitor cells. Eur J Neurosci. 2016;44(10):2858–70.

49. Erta M, Quintana A, Hidalgo J. Interleukin-6, a major cytokine in the central nervous system. Int J Biol Sci. 2012;8(9):1254–66.

50. Araujo DM, Cotman CW. Trophic effects of interleukin-4, -7 and -8 on hippocampal neuronal cultures: potential involvement of glial-derived factors. Brain Res. 1993 Jan;600(1):49–55.

51. Chistyakov D V., Astakhova AA, Azbukina N V., Goriainov S V., Chistyakov V V., Sergeeva MG. Cellular model of endotoxin tolerance in astrocytes: Role of interleukin 10 and oxylipins. Cells. 2019;8(12):1–10.

52. Guryleva M V., Chistyakov D V., Lopachev A V., Goriainov S V., Astakhova AA, Timoshina YA, et al. Modulation of the primary astrocyte-enriched cultures’ oxylipin profiles reduces neurotoxicity. Metabolites. 2021;11(8).

53. Bhatt S, Puli L, Patil CR. Role of reactive oxygen species in the progression of Alzheimer ‘ s disease. Drug Discov Today. 2021;26(3):794–803.

54. Firuzi O, Zhuo J, Chinnici CM, Wisniewski T, Praticò D. 5-Lipoxygenase gene disruption reduces amyloid-beta pathology in a mouse model of Alzheimer’s disease. FASEB J Off Publ Fed Am Soc Exp Biol. 2008 Apr;22(4):1169–78.

55. Manev H, Chen H, Dzitoyeva S, Manev R. Cyclooxygenases and 5-lipoxygenase in Alzheimer’s disease. Prog Neuropsychopharmacol Biol Psychiatry. 2011;35(2):315–9.

56. Kahnt AS, Angioni C, Göbel T, Hofmann B, Roos J, Steinbrink SD, et al. Inhibitors of Human 5-Lipoxygenase Potently Interfere With Prostaglandin Transport. 2022;12(January):1–13.

57. Steinhilber D. 5-Lipoxygenase: a target for antiinflammatory drugs revisited. Curr Med Chem. 1999 Jan;6(1):71–85.

58. Lin HC, Lin TH, Wu MY, Chiu YC, Tang CH, Hour MJ, et al. 5-lipoxygenase inhibitors attenuate TNF-α-Induced inflammation in human synovial fibroblasts. PLoS One. 2014;9(9).

59. Lu W, Maheshwari A, Misiuta I, Fox SE, Chen N, Zigova T, et al. Neutrophil-specific chemokines are produced by astrocytic cells but not by neuronal cells. Brain Res Dev Brain Res. 2005 Mar;155(2):127–34.

60. Kamphuis W, Kooijman L, Orre M, Stassen O, Pekny M, Hol EM. GFAP and vimentin deficiency alters gene expression in astrocytes and microglia in wild-type mice and changes the transcriptional response of reactive glia in mouse model for Alzheimer’s disease. Glia. 2015;63(6):1036–56.

61. Ashutosh, Kou W, Cotter R, Borgmann K, Wu L, Persidsky R, et al. CXCL8 protects human neurons from amyloid-β-induced neurotoxicity: Relevance to Alzheimer’s disease. Biochem Biophys Res Commun. 2011;412(4):565–71.

62. Walker DG, Lue LF, Beach TG. Gene expression profiling of amyloid beta peptide-stimulated human post-mortem brain microglia. Neurobiol Aging. 2001;22(6):957–66.

63. Galimberti D, Schoonenboom N, Scheltens P, Fenoglio C, Bouwman F, Venturelli E, et al. Intrathecal chemokine synthesis in mild cognitive impairment and Alzheimer disease. Arch Neurol. 2006;63(4):538–43.

64. Craig-Schapiro R, Kuhn M, Xiong C, Pickering EH, Liu J, Misko TP, et al. Multiplexed immunoassay panel identifies novel CSF biomarkers for alzheimer’s disease diagnosis and prognosis. PLoS One. 2011;6(4).

65. Kamphuis W, Middeldorp J, Kooijman L, Sluijs JA, Kooi EJ, Moeton M, et al. Glial fibrillary acidic protein isoform expression in plaque related astrogliosis in Alzheimer’s disease. Neurobiol Aging. 2014;35(3):492–510.

66. Bajetto A, Bonavia R, Barbero S, Schettini G. Characterization of chemokines and their receptors in the central nervous system: Physiopathological implications. J Neurochem. 2002;82(6):1311–29.

67. Persson T, Monsef N, Andersson P, Bjartell A, Malm J, Calafat J, et al. Expression of the neutrophil-activating CXC chemokine ENA-78/CXCL5 by human eosinophils. Clin Exp Allergy. 2003;33(4):531–7.

68. Xia M, Qin S, McNamara M, Mackay C, Hyman BT. Interleukin-8 receptor B immunoreactivity in brain and neuritic plaques of Alzheimer’s disease. Am J Pathol. 1997;150(4):1267–74.

69. Semple BD, Kossmann T, Morganti-Kossmann MC. Role of chemokines in CNS health and pathology: A focus on the CCL2/CCR2 and CXCL8/CXCR2 networks. J Cereb Blood Flow Metab. 2010;30(3):459–73.

70. Menten P, Wuyts A, Van Damme J. Monocyte chemotactic protein-3. Eur Cytokine Netw. 2001;12(4):554–60.

71. Renner NA, Ivey NS, Redmann RK, Lackner AA, MacLean AG. MCP-3/CCL7 production by astrocytes: implications for SIV neuroinvasion and AIDS encephalitis. J Neurovirol. 2011 Apr;17(2):146–52.

72. Thompson WL, Van Eldik LJ. Inflammatory cytokines stimulate the chemokines CCL2/MCP-1 and CCL7/MCP-3 through NFkB and MAPK dependent pathways in rat astrocytes [corrected]. Brain Res. 2009 Sep;1287:47—57.

73. Kumar A, Behl T, Jamwal S, Kaur I, Sood A, Kumar P. Exploring the molecular approach of COX and LOX in Alzheimer ‘ s and Parkinson ‘ s disorder. Mol Biol Rep. 2020;47(12):9895–912.

74. Schmelzer KR, Kubala L, Newman JW, Kim IH, Eiserich JP, Hammock BD. Soluble epoxide hydrolase is a therapeutic target for acute inflammation. Proc Natl Acad Sci U S A. 2005;102(28):9772–7.

75. Caggiano AO, Kraig RP. Prostaglandin E2 and 4-aminopyridine prevent the lipopolysaccharide-induced outwardly rectifying potassium current and interleukin-1beta production in cultured rat microglia. J Neurochem. 1998 Jun;70(6):2357–68.

76. Kalinski P. Regulation of Immune Responses by Prostaglandin E2. J Immunol. 2012;188(1):21–8.

77. Fattahi MJ, Mirshafiey A. Positive and negative effects of prostaglandins in Alzheimer’s disease. Psychiatry Clin Neurosci. 2014;68(1):50–60.

78. Casadesus G, Smith MA, Basu S, Hua J, Capobianco DE, Siedlak SL, et al. Increased isoprostane and prostaglandin are prominent in neurons in Alzheimer disease. Mol Neurodegener. 2007;2(1):1–8.

79. Basu S, Whiteman M, Mattey DL, Halliwell B, Ridge K, Haywood T. Raised levels of F 2 -isoprostanes and prostaglandin F 2 in di V erent rheumatic diseases. 2001;627–31.

80. Vane JR, Bakhle YS, Botting RM. Cyclooxygenases 1 and 2. Annu Rev Pharmacol Toxicol. 1998;38:97–120.

81. Nishizawa M, Nakajima T, Yasuda K, Kanzaki H, Sasaguri Y, Watanabe K, et al. Close kinship of human 20alpha-hydroxysteroid dehydrogenase gene with three aldoketo reductase genes. Genes Cells. 2000 Feb;5(2):111–25.

82. Arnaud L, Benech P, Greetham L, Stephan D, Jimenez A, Jullien N, et al. APOE4 drives inflammation in human astrocytes via TAGLN3 repression and NF-κB activation. Cell Rep. 2022;40(7).

83. Snow WM, Albensi BC. Neuronal gene targets of NF-κB and their dysregulation in alzheimer’s disease. Front Mol Neurosci. 2016;9(NOV2016):1–19.

84. Theendakara V, Peters-Libeu CA, Bredesen DE, Rao R V. Transcriptional Effects of ApoE4: Relevance to Alzheimer’s Disease. Mol Neurobiol. 2018;55(6):5243–54.

85. Theendakara V, Peters-Libeu CA, Spilman P, Poksay KS, Bredesen DE, Rao R V. Direct transcriptional effects of Apolipoprotein E. J Neurosci. 2016;36(3):685–700.

86. Gomez-Arboledas A, Davila JC, Sanchez-Mejias E, Navarro V, Nuñez-Diaz C, Sanchez-Varo R, et al. Phagocytic clearance of presynaptic dystrophies by reactive astrocytes in Alzheimer’s disease. Glia. 2018;66(3):637–53.

87. Konishi H, Koizumi S, Kiyama H. Phagocytic astrocytes: Emerging from the shadows of microglia. Glia. 2022;70(6):1009–26.

88. Schiweck J, Eickholt BJ, Murk K. Important ShapeshifterL: Mechanisms Allowing Astrocytes to Respond to the Changing Nervous System During Development, Injury and Disease. Front Cell Neurosci. 2018;12(August):1–17.

89. Tanaka K, Watase K, Manabe T, Yamada K, Watanabe M, Takahashi K, et al. Epilepsy and exacerbation of brain injury in mice lacking the glutamate transporter GLT-1. Science. 1997 Jun;276(5319):1699–702.

90. Rothstein JD, Martin L, Levey AI, Dykes-Hoberg M, Jin L, Wu D, et al. Localization of neuronal and glial glutamate transporters. Neuron. 1994 Sep;13(3):713–25.

91. Masliah E, Alford M, Mallory M, Rockenstein E, Moechars D, Van Leuven F. Abnormal glutamate transport function in mutant amyloid precursor protein transgenic mice. Exp Neurol. 2000 Jun;163(2):381–7.

92. Rosenberg PA, Aizenman E. Hundred-fold increase in neuronal vulnerability to glutamate toxicity in astrocyte-poor cultures of rat cerebral cortex. Neurosci Lett. 1989 Aug;103(2):162–8.

93. Diniz LP, Araujo APB, Matias I, Garcia MN, Barros-Aragão FGQ, de Melo Reis RA, et al. Astrocyte glutamate transporters are increased in an early sporadic model of synucleinopathy. Neurochem Int. 2020;138:104758.

94. Zelenaia O, Schlag BD, Gochenauer GE, Ganel R, Song W, Beesley JS, et al. Epidermal Growth Factor Receptor Agonists Increase Expression of Glutamate Transporter GLT-1 in Astrocytes through Pathways Dependent on Phosphatidylinositol 3-Kinase and Transcription Factor NF-κB. Mol Pharmacol. 2000 Apr 1;57(4):667LP–678.

95. Sitcheran R, Gupta P, Fisher PB, Baldwin AS. Positive and negative regulation of EAAT2 by NF-κB: A role for N-myc in TNFα-controlled repression. EMBO J. 2005;24(3):510–20.

96. Ghosh M, Yang Y, Rothstein JD, Robinson MB. Nuclear factor-κB contributes to neuron-dependent induction of glutamate transporter-1 expression in astrocytes. J Neurosci. 2011;31(25):9159–69.

97. Lee SG, Su ZZ, Emdad L, Gupta P, Sarkar D, Borjabad A, et al. Mechanism of ceftriaxone induction of excitatory amino acid transporter-2 expression and glutamate uptake in primary human astrocytes. J Biol Chem. 2008;283(19):13116–23.

98. Rodriguez-Kern A, Gegelashvili M, Schousboe A, Zhang J, Sung L, Gegelashvili G. Beta-amyloid and brain-derived neurotrophic factor, BDNF, up-regulate the expression of glutamate transporter GLT-1/EAAT2 via different signaling pathways utilizing transcription factor NF-kappaB. Neurochem Int. 2003;43(4–5):363–70.

99. Dresselhaus EC, Meffert MK. Cellular specificity of NF-κB function in the nervous system. Front Immunol. 2019;10(MAY).

100. Shi Y, Kirwan P, Smith J, Robinson H., Livesey FJ. Human cerebral cortex development from pluripotent stem cells to functional excitatory synapses. Nat Neurosci. 2012;15(3):477–86.

101. Serio A, Bilican B, Barmada SJ, Ando DM, Zhao C, Siller R, et al. Astrocyte pathology and the absence of non-cell autonomy in an induced pluripotent stem cell model of TDP-43 proteinopathy. Proc Natl Acad Sci U S A. 2013;110(12):4697–702.

102. Misheva M, Kotzamanis K, Davies LC, Tyrrell VJ, Rodrigues PRS, Benavides GA, et al. Oxylipin metabolism is controlled by mitochondrial β-oxidation during bacterial inflammation. Nat Commun. 2022;13(1):1–20.

103. Ostermann AI, Willenberg I, Schebb NH. Comparison of sample preparation methods for the quantitative analysis of eicosanoids and other oxylipins in plasma by means of LC-MS/MS. Anal Bioanal Chem. 2015 Feb;407(5):1403–14.

