## Supplemental data for "Upregulated NF-κB pathway proteins may underlie *APOE44* associated astrocyte phenotypes in sporadic Alzheimer’s disease"

**SUPPLEMENTARY DATA**

S1: PRIMERS USED FOR qPCR

| Gene | Marker | Forward Sequence | Reverse Sequence |
|---|---|---|---|
| B-actin | Reference gene | TCACCACCACGGCCGAGCG | TCTCCTTCTGCATCCTGTCG |
| OCT3/4 | Stem cell | TTCTGGCGCCGGTTACAGAACCA | GACAACAATGAAAATCTTCAGGAGA |
| Nanog | Stem cell | GCTTGCCTTGCTTTGAAGCA | TTCTTGACCGGGACCTTGTC |
| SOX2 | Stem cell/NPC | ACTTTTGTCGGAGACGGAGA | GTTCATGTGCGCGTAACTGT |
| PAX6 | NPC | GTGTCCAACGGATGTGTGAG | CTAGCCAGGTTGCGAAGAAC |
| FOXG1 | NPC | AGGAGGGCGAGAAGAAGAAC | TCACGAAGCACTTGTTGAGG |
| MAP2 | Neurons | AACCGAGGAAGCATTGATTG | TTCGTTGTGTCGTGTTCTCA |
| Vimentin | Astrocyte | TGGACCAGCTAACCAACGAC | GCCAGAGACGCATTGTCAAC |
| GFAP | Astrocyte | GAGCAGGAGGAGCGGCAC | TAGTCGTTGGCTTCGTGCTT |
| S100b | Astrocyte | ATGTCTGAGCTGGAGAAGGC | TTCAAAGAACTCGTGGCAGG |
| SLC1A3 (EAAT1) | Astrocyte | CGAAGCCATCATGAGACTGGTA | TCCCAGCAATCAGGAAGAGAA |
| SLC1A3 (EAAT2) | Astrocyte | TAACTCTGGCGGCCAATGGAAAGT | ACGCTGGGGAGTTTATTCAAGAAT |
| GLUL | Astrocyte | CCTGCTTGTATGCTGGAGTC | GATCTCCCATGCTGATTCCT |
| CD44 | Astrocyte precursor | AGCATCGGATTTGAGACCTG | GTTGTTTGCTGCACAGATGG |
| APOE | Apo-lipoprotein | GAGCAGGCCCAGCAGATAC | CTGCATGTCTTCCACCAGGG |
| C3 | Complement | AAAAGGGGCGCAACAAGTTC | GATGCCTTCCGGGTTCTCAA |
| MEGF10 | Phagocytosis | AGCGGATTTCAGAGAGGTTGT | GGTGATGCTGTCCCAATCCA |
| MERTK | Phagocytosis | TGGACAACTCTTCACAGCGG | GCAGAGGAATATGCTTTGGTCC |

S2: ANTIBODIES USED FOR IMMUNOCYTOCHEMISTRY

| Target | Marker of | Species | Dilution | Supplier | Cat no |
|---|---|---|---|---|---|
| OCT3/4 | pluripotent stem cells | Goat | 1:500 | Santa Cruz | SC-8628 |
| Tra-1-60 | pluripotent stem cells | Mouse | 1:100 | Santa Cruz | SC-21705 |
| SOX2 | pluripotent stem cells | Rabbit | 1:200 | Abcam | Ab97959 |
| Nestin | neuroectodermal stem cells | Mouse | 1:300 | BD Pharmigen | BD 611659 |
| FOXG1 | NPC | Rabbit | 1:250 | Abcam | ab18259 |
| PAX6 | NPC | Mouse | 1:1000 | DSHB | |

| | | | | | |
|---|---|---|---|---|---|
| PAX6 | NPC | Rabbit | 1:500 | Abcam | ab195045 |
| S100B | astrocytes | Mouse | 1:1000 | Sigma | S2532 |
| GFAP | astrocytes | Chicken | 1:1000 | Abcam | ab4674 |
| EAAT1 | astrocytes | Rabbit | 1:200 | Abcam | ab416 |

S3: ANTIBODIES USED FOR WESTERN BLOTTING

| Target | Marker of | Species | Dilution | Supplier | Cat no |
|---|---|---|---|---|---|
| GFAP | intermediate filament protein | Chicken | 1:1000 | Abcam | ab4674 |
| EAAT1 | glutamate receptor | Rabbit | 1:500 | Cell Signaling Technology | 5684 |
| EAAT2 | glutamate receptor | Mouse | 1:400 | Santa Cruz | sc-365634 |
| p65 | NF-κB protein | Mouse | 1:1000 | Cell Signaling Technology | 6956 |
| phospho-p65 | NF-κB protein | Rabbit | 1:500 | Cell Signaling Technology | 3033 |
| p105/50 | NF-κB protein | Rabbit | 1:1000 | Abcam | ab32360 |
| IκBα | NF-κB protein | Mouse | 1:500 | Cell Signaling Technology | 4814 |

S4: LIPIDS MEASURED USING LC/MS

| Lipid | Precursor m/z value | Ion m/z value |
|---|---|---|
| PGE2 and PGD2 | 351.2 | 271.1 |
| 15-deoxy-PGJ2 | 315.2 | 271.1 |
| TXB2 | 369.2 | 169.1 |
| 5-HETE | 319.2 | 115.1 |
| 8-HETE | 319.2 | 155.101 |
| 9-HETE | 319.2 | 167.1 |
| 11-HETE | 319.2 | 167.102 |
| 12-HETE | 319.2 | 179.1 |
| 15- HETE | 319.2 | 219 |
| 5-HEPE | 317.2 | 115.1 |
| 8-HEPE | 317.2 | 155.1 |
| 9-HEPE | 317.2 | 167.1 |
| 11-HEPE | 317.2 | 167.101 |
| 12-HEPE | 317.2 | 179.1 |
| 15-HEPE | 317.2 | 219.1 |
| 18-HEPE | 317.2 | 259.1 |
| 4-HDOHE | 343.2 | 101.1 |
| 7-HDOHE | 343.2 | 141.1 |
| 8-HDOHE | 343.2 | 189.1 |
| 10-HDOHE | 343.2 | 133.101, |
| 11-HDOHE | 343.2 | 121.1 |

| | | |
|---|---|---|
| 13-HDOHE | 343.2 | 193.1 |
| 14-HDOHE | 343.2 | 205.1 |
| 16-HDOHE | 343.2 | 233.101 |
| 20-HDOHE | 343.2 | 241.101 |
| 9-HODE | 295.2 | 171.1 |
| 13-HODE | 295.2 | 195.1 |
| 5-HETrE | 321.2 | 115.1 |
| 15-HETrE | 321.2 | 221.1 |
| 9,10-DiHOME | 313.2 | 201.1 |
| 12,13-DiHOME | 313.2 | 183.1 |
| 8,9-DiHETrE | 337.2 | 127.1 |
| 11,12-DiHETrE | 337.2 | 167.1 |
| 14,15-DiHETrE | 337.2 | 207.1 |
| 14,15-DiHETE | 335.201 | 207.1 |
| 17,18-DiHETE | 335.2 | 247.1 |

### S5: FULL RANGE OF CYTOKINES/CHEMOKINES DETECTED IN QUIESCENT ASTROCYTES

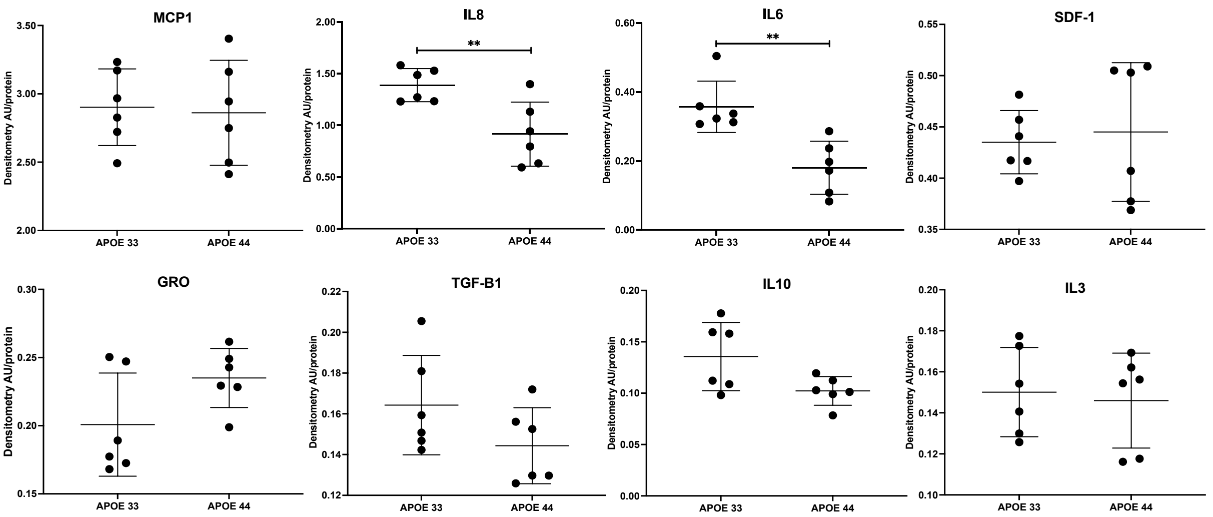


### S6: FULL RANGE OF CYTOKINES/CHEMOKINES DETECTED IN A1 ASTROCYTES

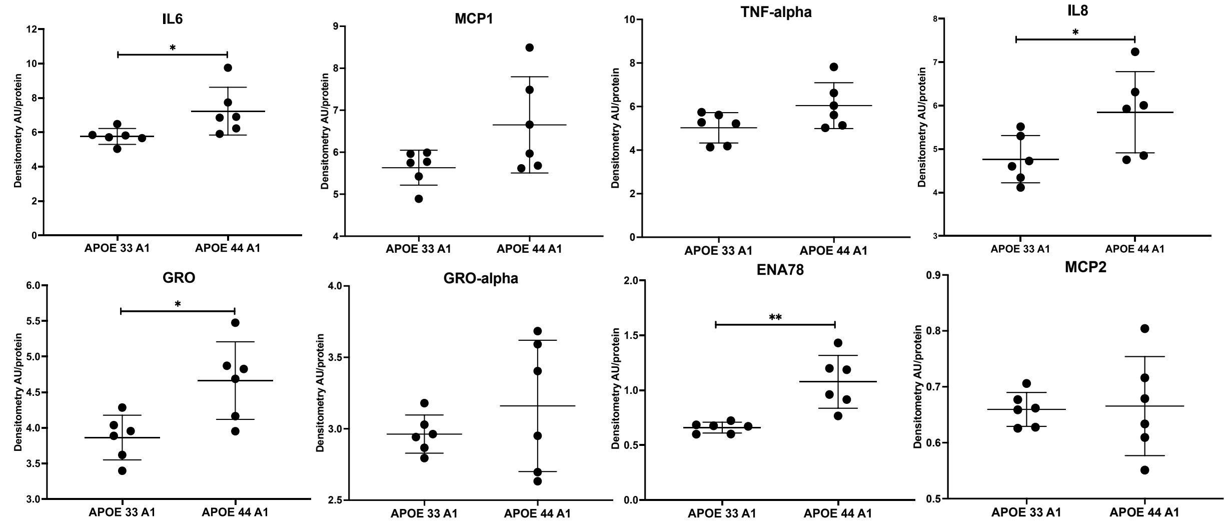

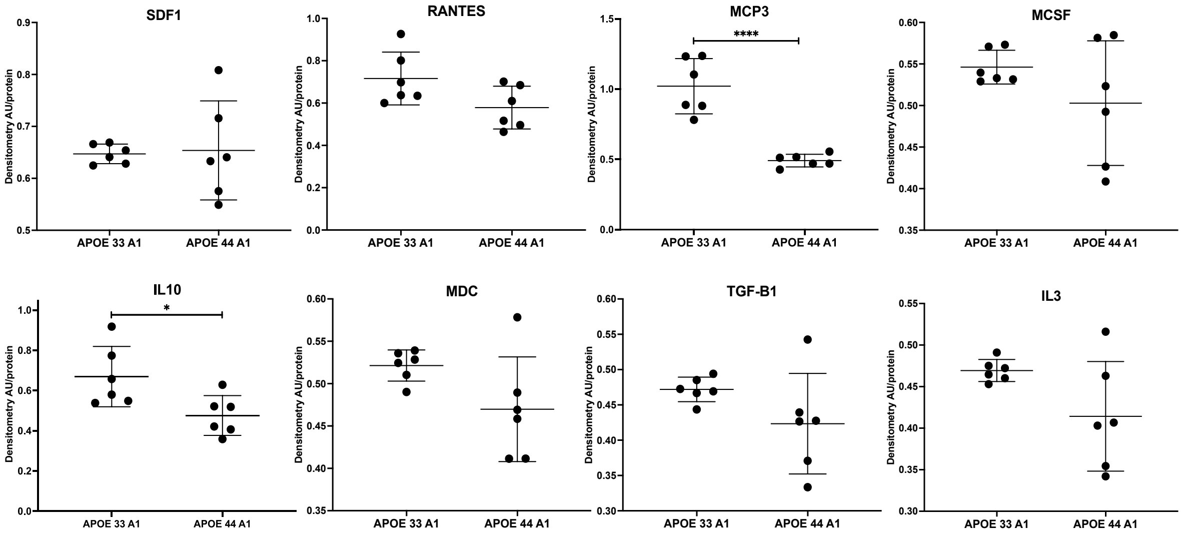


### S7: FULL RANGE OF OXYLIPINS DETECTED (QUIESCENT AND A1 SAMPLES)

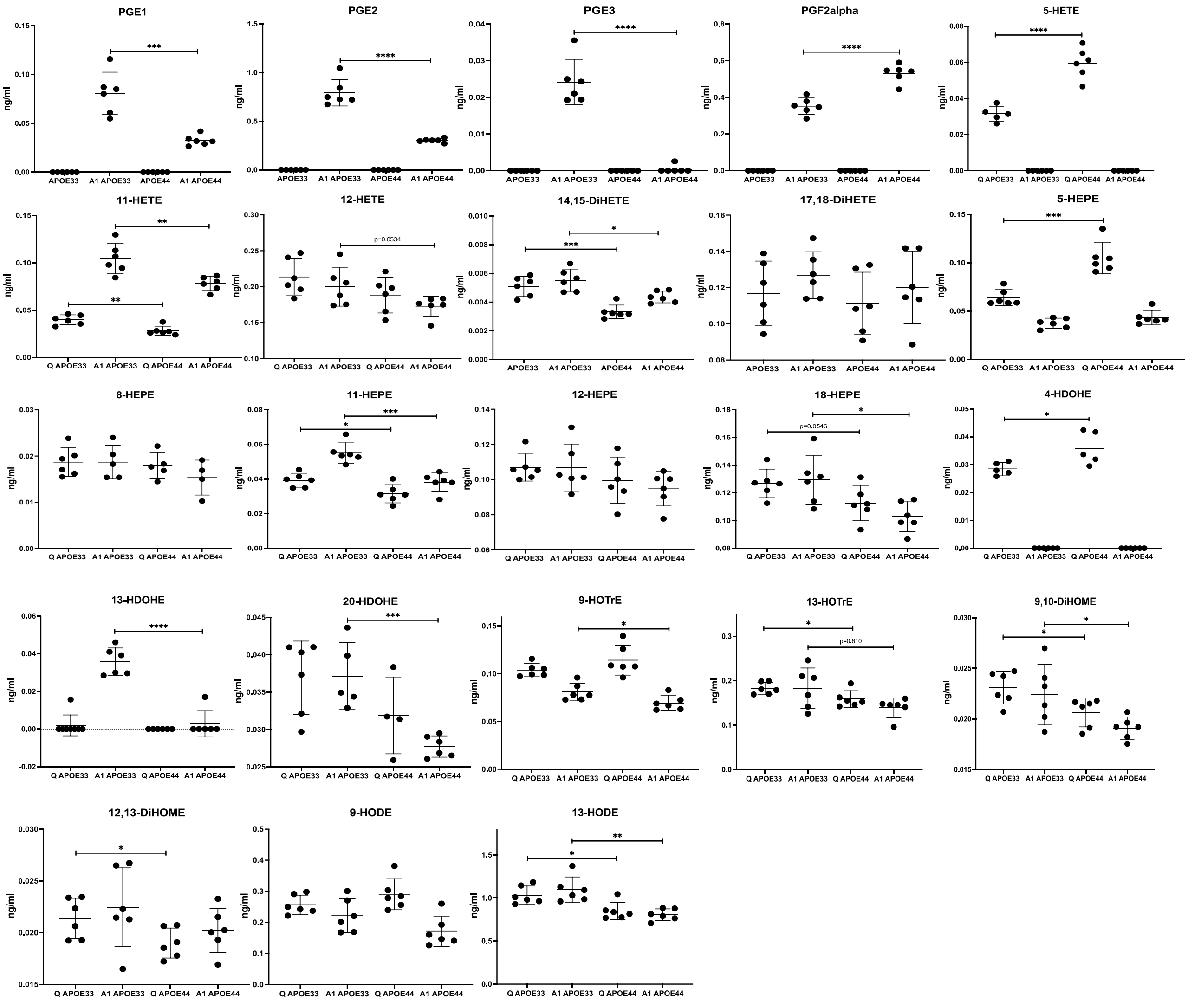
